# Extracellular Vacuole-derived bodies (EVacs) mediate RNA secretion in plants

**DOI:** 10.64898/2026.08.31.748268

**Authors:** M. Lucía Borniego, Meenu Singla-Rastogi, Megha H. Sampangi-Ramaiah, Ang-Yu Liu, Giovanni Gonzalez-Gutierrez, Sarah J. Cox-Vázquez, Chi-Tam Vo, Akihito Fukudome, Ricardo Javier Vázquez, Blake C. Meyers, Patricia Baldrich, Roger W. Innes

## Abstract

Extracellular RNAs are found in the plant extracellular space, but how they are exported from cells remains unclear. We found that the plant vacuole is a major source of extracellular RNA and identified a class of large <u>e</u>xtracellular <u>vac</u>uole-derived bodies, which we termed EVacs, that are key mediators of this transport. EVacs are marked by the vacuolar membrane (tonoplast) proteins γ-TIP and V-ATPase and originate as intravacuolar structures formed by inward folding of the tonoplast, encapsulating intact cytoplasmic material, including both RNAs and proteins. These intravacuolar bodies then escape the vacuole and are subsequently released from the plasma membrane of mesophyll cells into the apoplast. These findings provide a novel mechanism for the unconventional secretion of macromolecules in plants.

## Background

The exchange of molecular information between cells and across kingdoms is critical for plant adaptation and defense. Extracellular RNA (exRNA) has emerged as a key mediator of this communication (*1, 2*). To date, research in this field has focused on small RNAs associated with extracellular vesicles (EVs; 50-200 nm in diameter) (*3*). However, the majority of exRNA present in plant leaves is located outside of vesicles (*4, 5*). While some of these RNAs are stabilized by RNA-binding proteins or calcium-dependent condensates, they remain largely susceptible to extracellular ribonucleases (RNases) (*5, 6*). Consequently, exRNAs found inside leaves typically exhibit a fragmented profile characterized by transfer RNA-derived fragments (tRFs) and ribosomal RNA-derived fragments (rRFs), with little, if any, intact messenger RNA (mRNA) (*6*).

The mechanisms governing the secretion of extravesicular RNAs have remained essentially unexplored. While investigating the functional role of plant RNases in extravesicular RNA processing, we observed an unexpected correlation between RNA found inside plant vacuoles and exRNA. An *Arabidopsis* mutant deficient in vacuolar RNA catabolism accumulated long RNAs not only within the vacuole but also in the extracellular space of plant leaves, which is referred to as the apoplast. This unexpected correlation, along with the detection of integral vacuolar membrane (tonoplast) proteins in the apoplast, suggested a direct physical communication between these two compartments that had remained overlooked. In this study, we demonstrate that the vacuole is a major source of apoplastic RNA and identify a novel class of large, <u>e</u>xtracellular <u>vac</u>uole-derived bodies, which we termed EVacs, that mediate this transport. We show that EVacs encapsulate un-processed RNA species, including full-length tRNAs, and 5.8S, and 5S rRNAs that are actively secreted into the apoplast. Upon secretion, EVacs likely undergo lipolytic disintegration mediated by apoplastic lipases, releasing their macromolecular cargo, and exposing full-length RNAs to apoplastic RNases.

### Extracellular RNAs are processed by pH-dependent apoplastic RNases

Among the diverse RNases secreted by plants, members of the T2 family are the most well-characterized and are known to be secreted into the apoplast (*7, 8*). To investigate the extent to which RNase-mediated processing shapes the apoplastic RNA pool, we assessed the ability of apoplastic wash fluid (AWF) to degrade total cell lysate (CL) RNA. AWF degraded most of it in just minutes (fig. S1A), highlighting the apoplast as a major compartment for RNA processing. Consistent with this RNA degradation activity, immunoblot analysis revealed that AWF contains the extracellular T2 RNase RNS3 (fig. S1B).

Since apoplastic RNAs are exposed to RNase processing, we assessed exRNA profiles under reduced RNase activity. Given the lack of effective inhibitors for plant T2-type RNases, and their known acidic pH optima (*9*), we isolated AWF using a range of high pH buffers. Alkalinization of the apoplast shifted the RNA profile toward increased abundance and longer species (Fig. 1A). Notably, the AWF profile at pH 8.2 closely resembled total cell lysate (CL) RNA, indicating a near-complete inhibition of apoplastic RNase activity. This was further confirmed by exposing total CL RNA to AWF at pH 8.2 (Fig. 1B). Even a moderate shift to pH 7.3 was sufficient to significantly reduce degradation in the AWF. Under these conditions, the AWF RNA was processed slowly, yielding the characteristic AWF RNA profile only after a 2-hour incubation at room temperature (Fig. 1C).

**Fig. 1.**
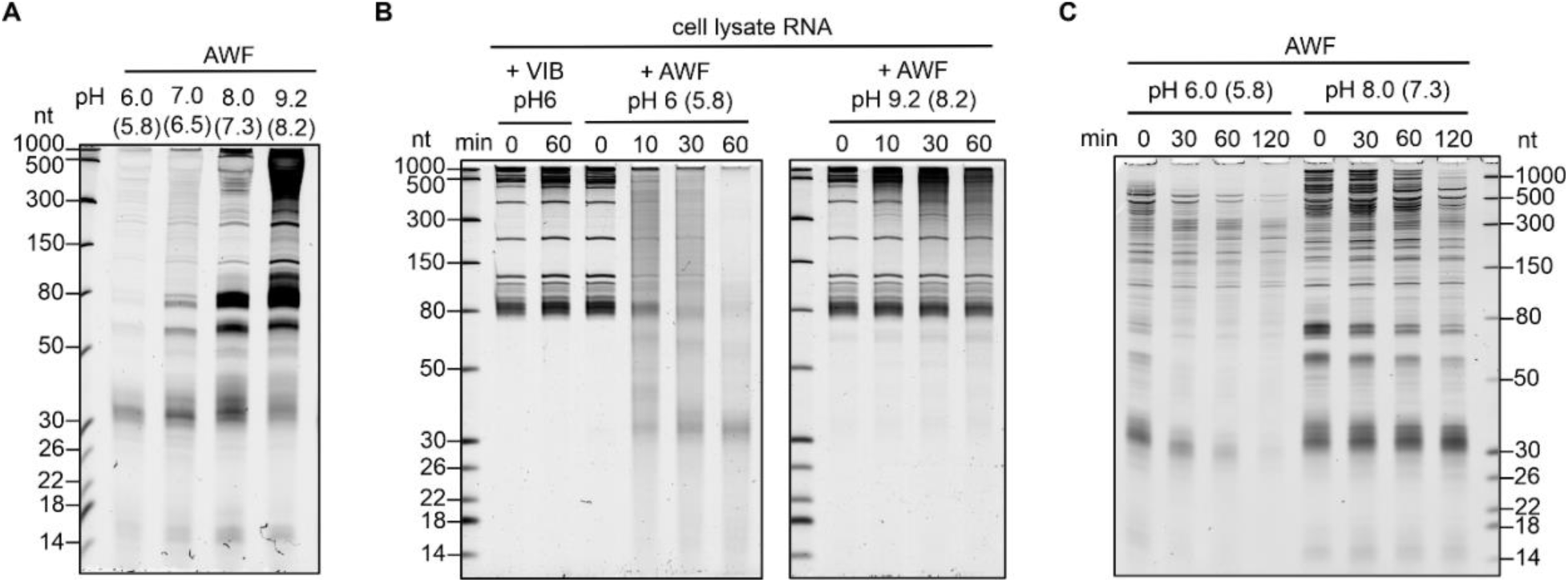
Post-secretion processing by apoplastic RNases shapes the extracellular RNA pool in a pH-dependent manner. **(A)** AWF was isolated using buffers of increasing pH to reduce RNase activity. RNA was extracted from equal volumes and resolved by denaturing polyacrylamide gel electrophoresis (PAGE). The indicated values are the pH values of the buffers used for AWF extraction; bracketed values indicate the actual pH values measured in the collected AWF samples. Ladder sizes in nucleotides (nt). **(B)** Total cell lysate RNA was incubated at RT for the indicated times with Vesicle Isolation Buffer (VIB) or AWF isolated using different pH buffers for the indicated times prior to TRIzol extraction and denaturing PAGE. Values indicate extraction buffer pH; bracketed values indicate pH measured in collected AWF. **(C)** AWF isolated at the indicated pH values was incubated at RT for the indicated times prior to TRIzol extraction and denaturing PAGE.

Although RNases belonging to the T2 family are the primary enzymes implicated in plant tRNA processing (*10*), the specific contribution of the apoplastic members RNS1 and RNS3 remains poorly defined. Our analysis of AWF from *rns1*, *rns3*, and *rns1/rns3* double mutants revealed no significant reduction in RNA-processing capacity compared to wild-type levels (fig. S2). These results demonstrate a high degree of functional redundancy among apoplastic RNases and suggest that the exRNA landscape is shaped by a complex, multi-enzymatic network.

### The vacuole is a major source of apoplastic RNA

We next investigated whether intracellular RNases contribute to the exRNA landscape. RNS2 is the primary RNase responsible for RNA decay within the vacuole (*9, 11*). Its absence triggers constitutive autophagy, leading to the accumulation of long RNAs, predominantly rRNA, within the vacuole (*11, 12*). Analysis of RNA isolated from wild-type vacuoles showed that vacuolar RNA is less degraded than AWF RNA (fig. S3A). This observation suggested that the vacuole could be a source of exRNA. The *rns2-2* mutant showed a large increase in vacuolar RNA, including rRNAs and tRNAs, when normalized by leaf fresh weight (fig. S3A). When we normalized gel loading by RNA amount, we observed a higher accumulation of both sRNAs and RNAs longer than 500 nt in *rns2-2* vacuoles (fig. S3B), indicating that RNS2 contributes to general degradation of RNA.

Notably, *rns2-2* AWF accumulated more intact rRNA species (fig. S4A) and slightly truncated tRNAs and tRFs, mirroring the vacuolar pool. Although RNS2 is detectable in the apoplastic proteome (*13*), the accumulation of these RNA species in the AWF of *rns2-2* does not seem to be a result of less extracellular RNase activity as *rns2-2* AWF retained full degradative activity (fig. S4B). Since the lack of RNS2 leads to increased autophagy, we addressed whether autophagy is involved in RNA secretion. Secretory autophagy has been reported in animals (*14*), but whether this also occurs in plants has not yet been confirmed (*15*). AWF RNA from an *atg5* mutant showed increased accumulation of tRNAs and tRFs; however, this accumulation was abolished in the *sid2/atg5* double mutant (fig. S5), indicating that it arises from elevated salicylic acid (SA) rather than loss of autophagy per se (*16*). These results show that autophagy is not a major pathway involved in RNA secretion and motivated us to further explore the vacuole as a source of exRNA.

### Large vacuole-derived extracellular bodies are secreted into the apoplast

Current models of vacuolar discharge into the apoplast primarily describe a pathogen-induced fusion between the tonoplast and the plasma membrane (PM). This mechanism facilitates the targeted release of vacuolar antibacterial proteins and is associated with programmed cell death (*17, 18*). Whether a similar process operates under basal conditions remains unknown. To explore this, we compared published vacuolar and apoplastic proteomes and found a significant overlap (fig. S6A). Remarkably, many integral tonoplast (plant vacuolar membrane) proteins were also detected in the apoplast (fig. S6A). We therefore assessed whether the vacuolar cargo protein aleurain, as well as the tonoplast markers γ-TIP (also known as TIP1-1) and V-ATPase subunit E1 (VHA-E1) were found in the apoplast. Immunoblotting of AWF proteins confirmed that all three proteins are present in the apoplast (fig. S6B). Notably, the vacuolar fraction was depleted of cytoplasmic and apoplastic proteins (fig. S7).

Under the current model of vacuolar discharge, the fusion between the tonoplast and plasma membrane would result in the extracellular delivery of only soluble cargo. The observed enrichment of integral tonoplast proteins in the apoplast indicates that there may be other mechanisms for vacuolar cargo release into the extracellular space. To explore this, we performed live-cell imaging using the γ-TIP-GFP reporter line. In this line, GFP is fused to the C-terminus of γ-TIP, a vacuolar membrane aquaporin (*19*). Since both the N- and C-termini of γ-TIP are exposed to the cytosol, the GFP signal is not quenched by the low pH of the vacuole. In addition to tonoplast, γ-TIP-GFP marks intravacuolar bodies, commonly referred to as ‘bulbs’ (*20, 21*). Through confocal microscopy of freshly mounted *Arabidopsis* leaves, we observed large γ-TIP-GFP-containing spherical structures, 1-5 µm in diameter, budding from the plasma membrane of mesophyll cells (Fig. 2, fig. S8). These tonoplast-derived structures subsequently detach from the plasma membrane and are released into the apoplast (Fig. 2, fig. S8, Movies S1-S6). While the cell wall is traditionally viewed as a rigid barrier, ultrastructural evidence from specialized plant tissues suggests that plant cell wall can accommodate the passage of large membranous bodies while maintaining their structural integrity (*22*). Together, these observations demonstrate that large, extracellular vacuole-derived bodies are actively secreted into the extracellular environment. We hereafter refer to these structures as EVacs for <u>E</u>xtracellular <u>Vac</u>uole-derived bodies.

**Fig. 2.**
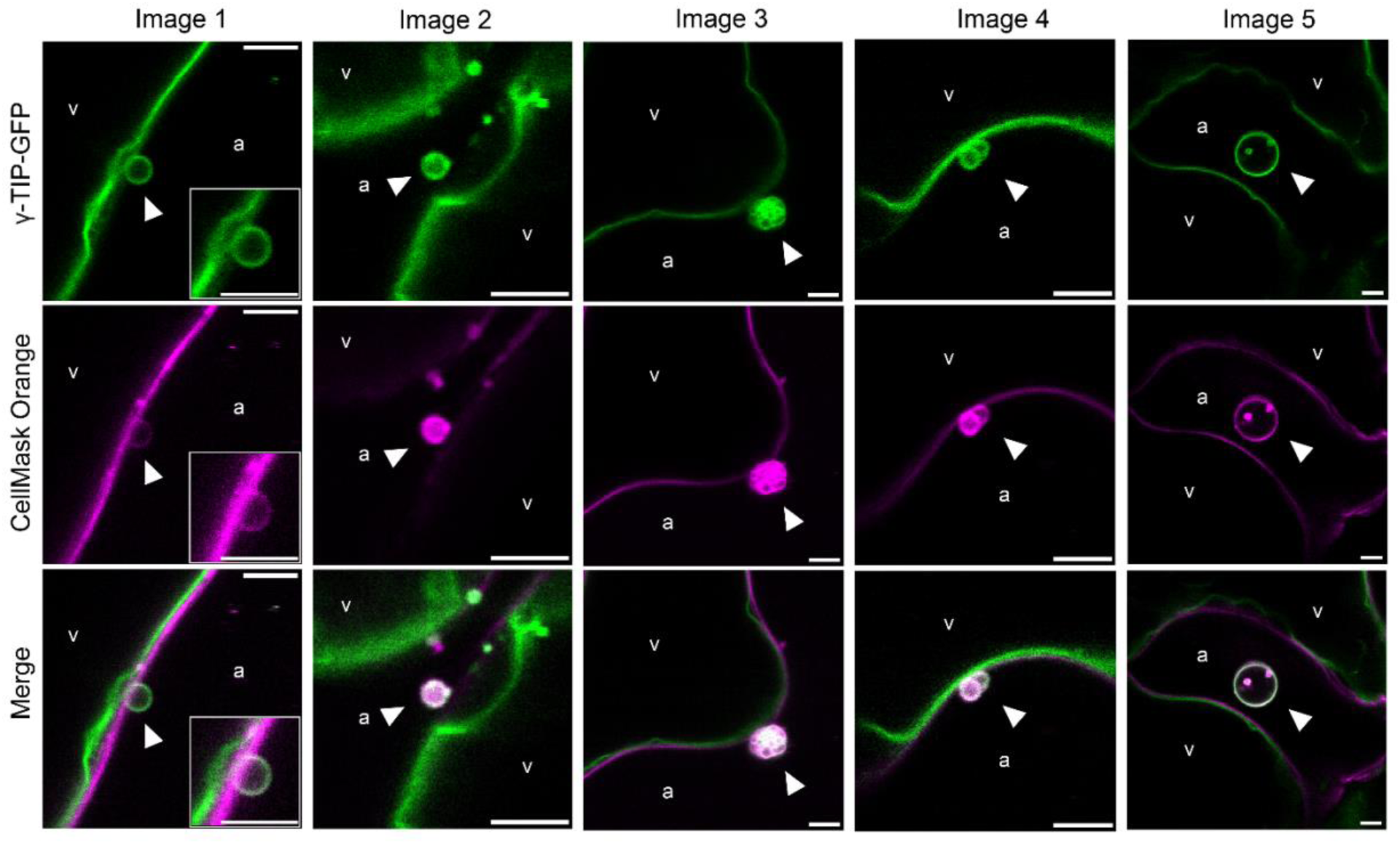
Extracellular vacuole-derived vesicles (EVacs) containing γ-TIP-GFP are released into the apoplast. In vivo confocal imaging of mesophyll cells from freshly detached leaves of 6-week-old γ-TIP-GFP *Arabidopsis* plants. Plasma membrane was stained with CellMask Orange. To enable imaging of the apoplastic space, leaves were infiltrated with VIB or distilled water and mounted in VIB buffer immediately prior to imaging. Images were acquired on a Leica Stellaris 8 confocal microscope. Vacuoles (v) and apoplast (a) are indicated. Arrowheads mark EVacs. The inset in image 1 shows a magnified view of the EVac region. Scale bars, 5 µm. See fig. S8 and Movies S1–S6 for additional examples.

### EVacs contain vacuolar proteins and are diverse in morphology

To characterize the molecular cargo of EVacs, we developed a gentle fractionation protocol using unfiltered AWF, as their large size and inherent fragility make them susceptible to mechanical disruption. Isolated EVacs appeared as spherical bodies under confocal microscopy; notably, many exhibited a distinct region of intense γ-TIP-GFP signal, possibly marking the site of abscission from the plasma membrane (Fig. 3A, fig. S9). Fractionation via differential centrifugation revealed that most of the γ-TIP-GFP signal concentrated in the low-speed pellet (2,000 *g*; P2), with a smaller population recovered at 15,000 *g* (P15-P2) (Fig. 3B, fig. S9). The detection of the tonoplast marker VHA-E1 in these same fractions further confirmed that EVacs are delimited by tonoplast membrane. Importantly, EVacs were also identified in wild-type plants (fig. S10), demonstrating that their secretion is a physiological process rather than an artifact of reporter overexpression.

**Fig. 3.**
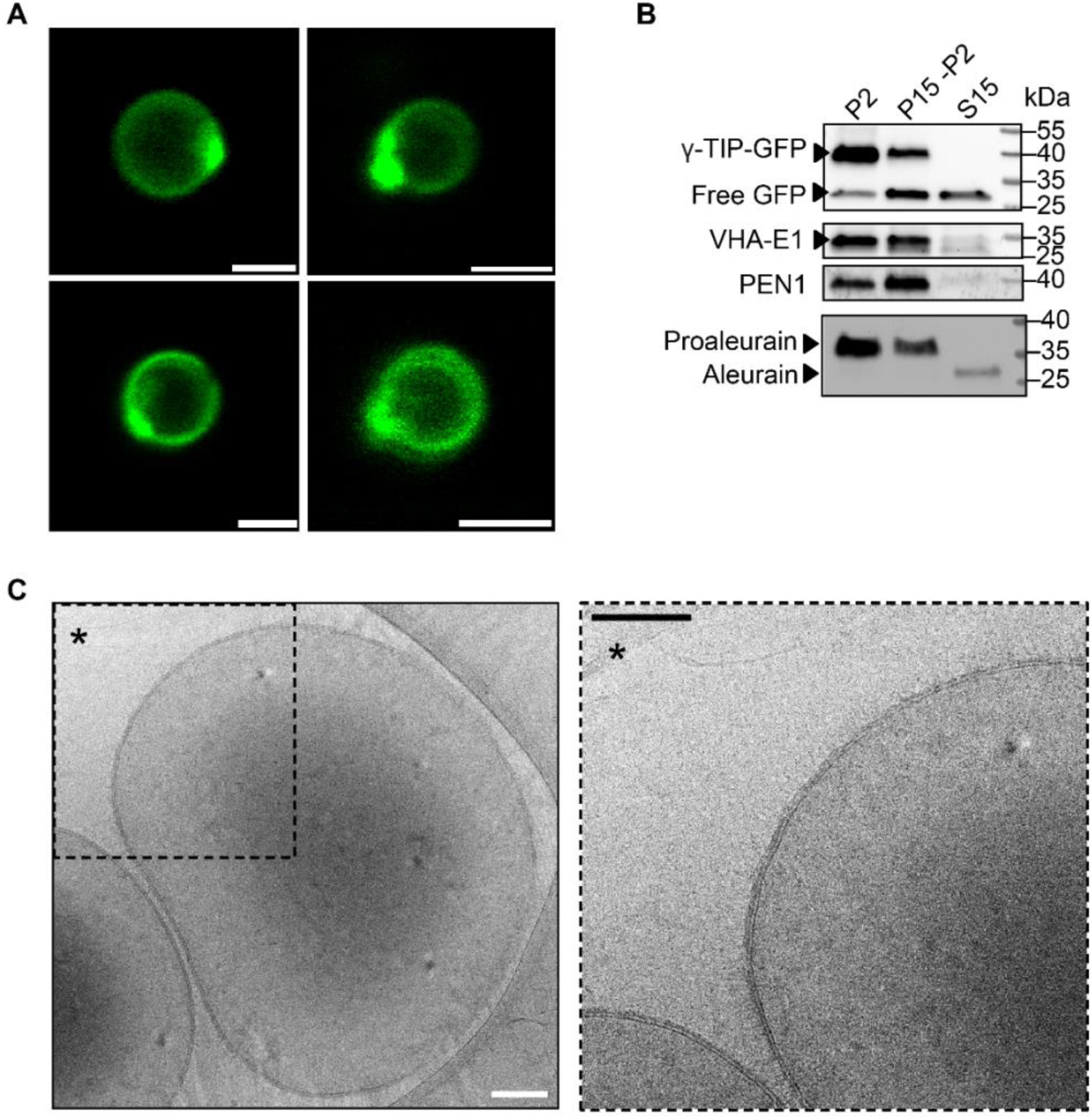
EVacs contain vacuolar proteins and are diverse in morphology. **(A)** Confocal images of EVacs (P2) isolated from unfiltered AWF of γ-TIP-GFP *Arabidopsis* leaves. A z-projection (top left) and single optical sections (top right and bottom) are shown. Scale bar, 2 µm. **(B)** Immunoblots detecting vacuolar markers in EVac-containing fractions (P2 and P15–P2) and the corresponding supernatant (S15). Loading is described in Materials and Methods. **(C)** Cryo-electron micrographs (cryo-EM) of purified EVacs. The right panel displays a high-magnification view of the dashed boxed region (*) from the overview image (left), showing a lipid bilayer surrounded by an external corona. Scale bar, 100 nm.

Consistently, proteomic analysis of isolated EVacs identified numerous vacuolar and tonoplast-resident proteins, including multiple subunits of the V-type proton ATPase complex, the pyrophosphatase proton pump AVP1, TIP1-1, and TIP1-2, alongside luminal hydrolases including the thiol protease aleurain (Data S1). To further characterize the luminal protein cargo of EVacs, we analyzed the vacuolar protease aleurain. Aleurain is synthesized as a ∼37 kDa pro-enzyme that is trafficked via prevacuolar compartments/multivesicular bodies (PVCs/MVBs) to the vacuole, where it undergoes proteolytic processing to its mature ∼28 kDa form (*23–27*).

While the mature form is the predominant species in the vacuole, whole AWF and supernatant fractions, the proenzyme is specifically enriched in the EVac fractions (Fig. 3B, fig. S6B, fig. S7, fig. S10B). This indicates that EVacs contain cargo that has progressed through the Golgi but has not yet been exposed to the degradative environment of the vacuolar lumen. Upon post-release rupture of EVacs, proaleurain likely undergoes extracellular maturation mediated by apoplastic acidic proteases, such as the RD21-like enzymes frequently identified in AWF proteomes (*13, 24*). These findings establish EVacs as a unique class of large extracellular organelles with a distinct protein signature and a protected internal environment.

Confocal imaging of mesophyll cells in intact cotyledons and 10,000 *g* (P10) fractions harvested from mature plants showed that EVacs can also encapsulate chloroplasts or chloroplast fragments (fig. S11A, B). This is also supported by our proteomic analysis, confirming an enrichment of chloroplast proteins in EVac fractions (Data S1). Notably, free chloroplasts not associated with EVacs were also detected in the P10 fractions. Although some may have been released upon EVac lysis in the apoplast, others might represent debris from natural cell turnover and death within the leaf tissue. Finally, we performed cryo-electron microscopy (cryo-EM) to resolve the EVac ultrastructure. Although some were multilayer, most of them were delimited by a single lipid bilayer that occasionally appeared double in specific regions, and many featured an external corona, presumably composed of glycoproteins (Fig. 3C). While technical limitations in vitreous ice thickness (∼100 nm) bias cryo-EM toward smaller EVacs, these images revealed a high degree of internal complexity, including electron-dense luminal content and, in some instances, intraluminal vesicles (fig. S12).

To distinguish EVacs from conventional EV secretion, we assessed the presence of the EV marker protein PENETRATION1 (PEN1), a syntaxin family protein, in EVacs isolated from a dual-labeled *RFP-PEN1 γ-TIP-GFP* line. Many EVacs contained RFP-PEN1 (Fig. 4A, Movie S7). Consistent with PEN1 being packaged into EVacs, immunoblots of P10 fractions isolated from the dual-labeled line revealed abundant PEN1 protein (Fig. 4B), showing a clear enrichment in the P10 fraction relative to the supernatant (fig. S13A). Like PEN1, the EV-marker TETRASPANIN8 (TET8) also occasionally compartmentalized inside EVacs, displaying a punctate or diffuse pattern. However, unlike PEN1, no evident colocalization of TET8 with the EVac membrane was observed (Fig. 4C). Like PEN1, TET8 was enriched in the P10 fraction relative to the supernatant (Fig. 4D, fig. S13B). The presence of EV-like particles within the EVac lumen, together with the detection of EV markers inside EVacs, raises the possibility that EVacs may themselves serve as vehicles for the secretion of EVs into the apoplast.

**Fig. 4.**
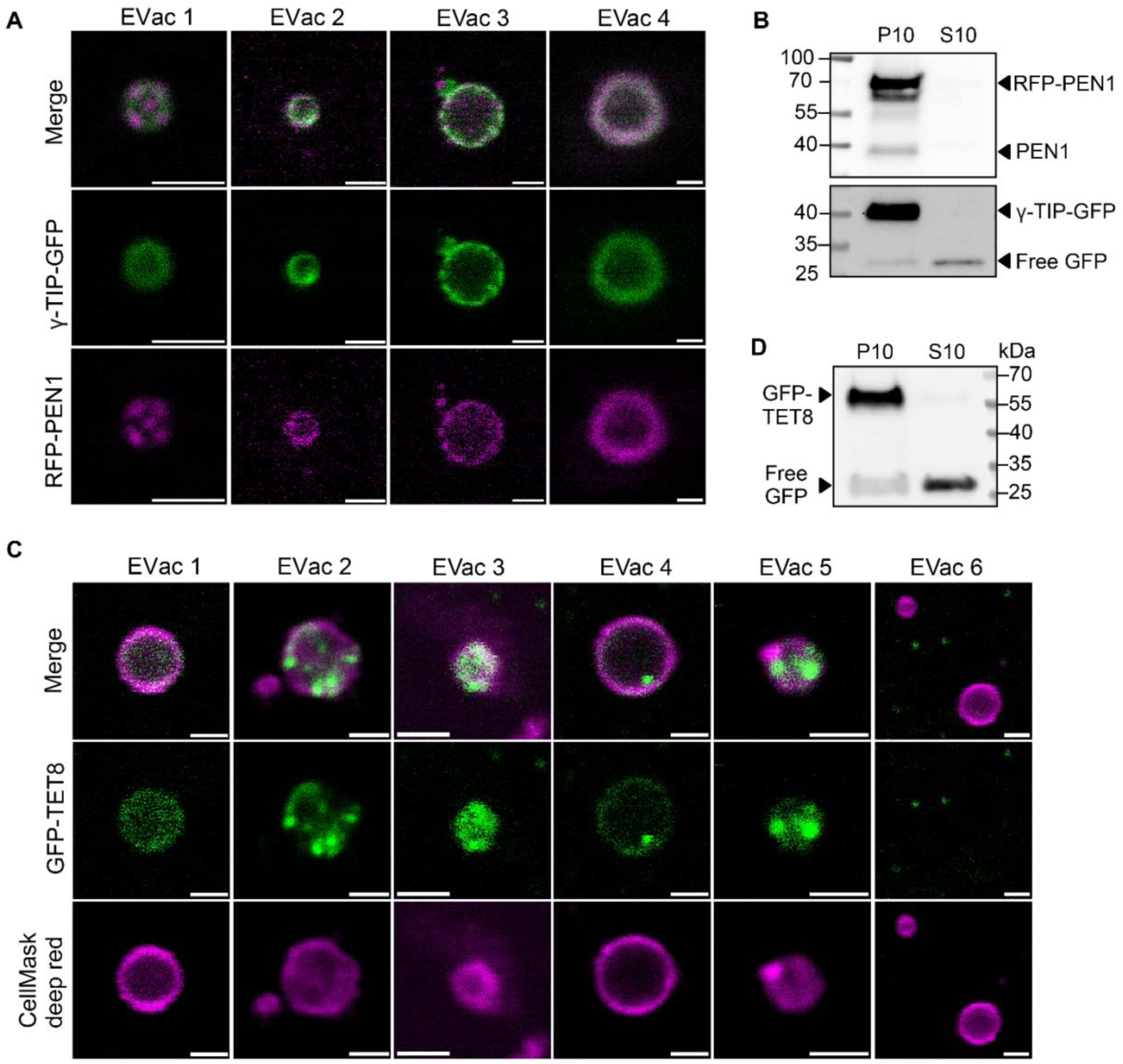
The extracellular vesicle markers PEN1 and TET8 localize to EVacs. **(A)** Confocal images of EVacs (P10) isolated from the γ-TIP-GFP × PEN1-RFP reporter line showing γ-TIP-GFP and PEN1-RFP localization at the EVac boundary membrane and within the EVac lumen. Scale bar, 2 μm. **(B)** Immunoblot detecting γ-TIP-GFP and PEN1-RFP in P10 and S10 fractions. Blot was probed with anti-GFP and anti-PEN1 antibodies. Loading is described in Materials and Methods. **(C)** Confocal images of EVacs (P10 fraction) isolated from the ProTET8::GFP-TET8 line. EVac membrane was labeled using CellMask deep red. Panels 1-6 show representative GFP-TET8 localization patterns, including the absence of GFP-TET8 signal in EVac 6. Scale bar, 2 μm. **(D)** Immunoblot detecting GFP-TET8 in P10 and S10 fractions. Blot was probed with an anti-GFP antibody. Loading is described in Materials and Methods.

### EVacs carry full-length tRNAs and rRNAs

Given that vacuoles accumulate diverse RNA species, we investigated whether EVacs transport RNA to the apoplast. As a first assessment, we labeled purified EVacs with the RNA-specific dye SYTO RNASelect, which revealed strong fluorescence inside EVacs, indicating they contained RNA (Fig. 5A). Analysis of RNA size distributions revealed that EVac preps contain RNAs ranging from ∼30 nt to >1,000 nt in length (Fig. 5B, fig. S14A). RNA gel-blot analysis of common extracellular tRNAs and rRNAs (*6*) further confirmed that EVacs contain full length tRNAs, and 5.8S and 5S rRNAs (Fig. 5C, fig. S14B). Notably, the smaller cleavage products (tRFs) typically generated by RNase activity are depleted in EVac preps, further supporting the non-degradative nature of EVacs. An RNase protection assay further confirmed the presence of full-length tRNAs and 5.8S/5S rRNAs inside the EVacs (Fig. 5D). tRNA fragments were enriched in the EVac-depleted supernatant fraction (Fig. 5C), representing the cargo released from EVacs upon their rupture in the apoplast, where the RNA is exposed to RNase activity. The EVac cargo release to the apoplast is likely driven by lipolytic degradation of the EVac membrane. Consistent with this, LC-MS/MS proteomic profiling of the P10 fraction showed a major enrichment of lipid-degrading machinery (Data S1). Notably, the GDSL esterase/lipase At1g29670 was identified with high confidence across replicates, supported by GDSL lipases At1g29660 and ESM1, alongside key phospholipases (PLD, PLC, and PLA2-α). This suggests that EVacs undergo controlled lipolytic disintegration upon secretion, systematically releasing their cargo into the apoplast.

**Fig. 5.**
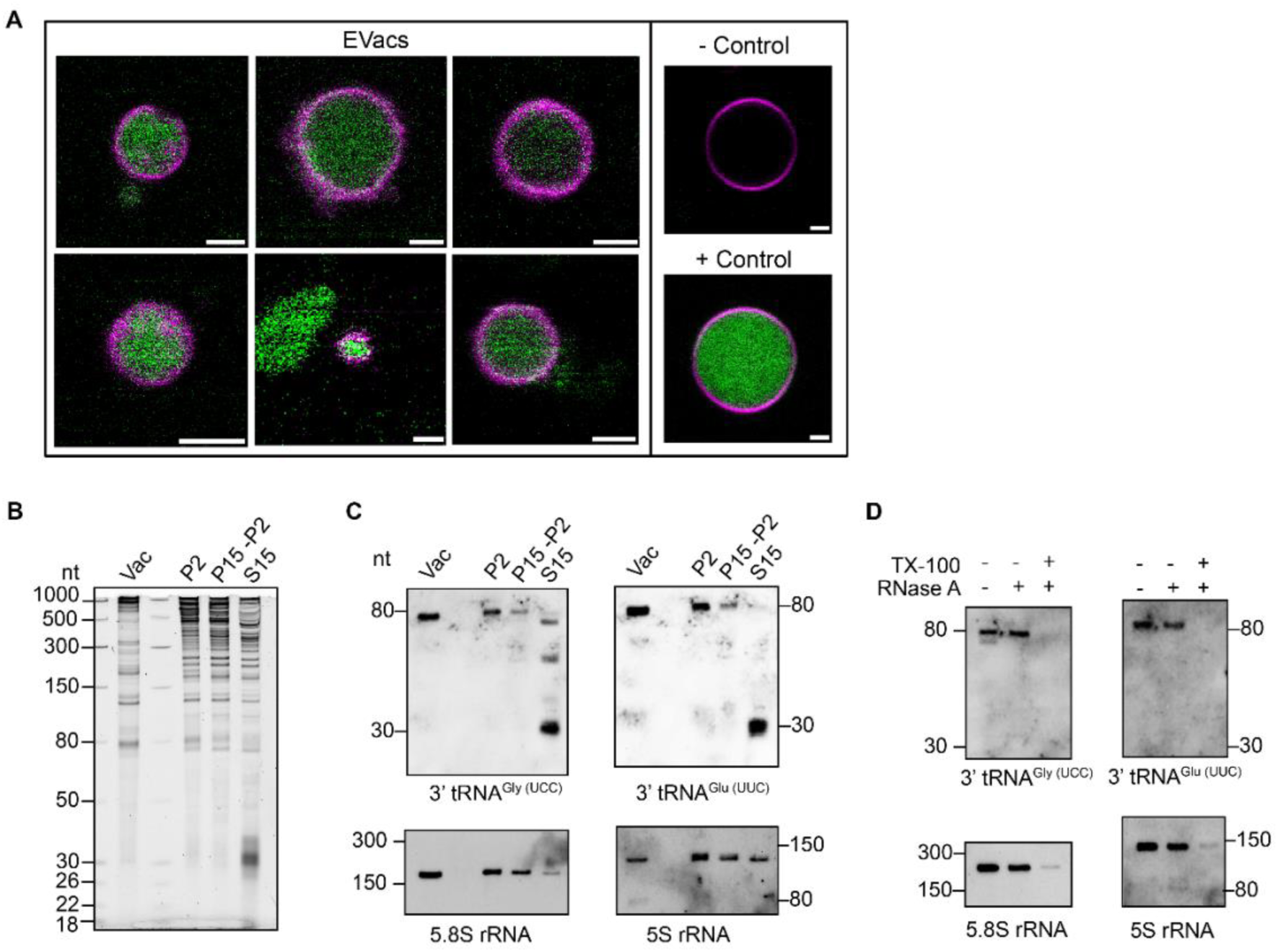
EVacs contain full-length RNA species. **(A)** Confocal images of γ-TIP-CFP EVacs (magenta) labeled with the RNA-specific dye SYTO RNASelect (green). γ-TIP-CFP was used to avoid spectral overlapping with the green RNA dye. Empty and RNA-loaded synthetic vesicles served as negative and positive controls, respectively. CellMask Deep red was used to label the synthetic vesicle membrane (in magenta). Scale bar, 2 µm. **(B)** RNA profile of vacuoles (Vac) and EVac fractions resolved by denaturing PAGE. Loading is described in Materials and Methods. **(C)** RNA gel-blot detection of the indicated RNAs from the gel shown in (B). **(D)** RNase A protection assay showing that full-length tRNAs and 5.8S and 5S rRNAs are protected within EVacs in the absence of detergent.

Notably, cryo-EM images showed small non-vesicular electron-dense particles resembling intact ribosomes both outside and inside EVacs (fig. S15A). Preliminary 2D class averaging of these particles consistently recovered two distinct particle-size classes, approximately 150 Å and 300 Å in diameter (fig. S15B). The larger class is compatible in size with mature ribosomes, although its morphology was not sufficiently distinctive for confident identification (fig. S15B). Moreover, proteomics along with immunoblot analysis confirmed the presence of ribosomal proteins within the EVac fraction (fig. S15C, Data S1). As ribosomes normally do not pellet at the speed used for EVac isolation, the ribosomes outside EVacs were likely released due to the inherent fragility of the EVac membrane during isolation.

The potential presence of intact ribosomes inside EVacs suggested that EVacs might carry intact mRNAs. We therefore assessed the mRNA content of EVacs using 3’ poly A RNAseq analysis (Data S2). This analysis revealed that EVacs contain diverse mRNAs that largely mirror the mRNA population found in whole cell lysate (fig. S16). The presence of full-length rRNAs, tRNAS and ribosomal proteins in EVacs suggests EVacs may sequester functional translation complexes. While autonomous extracellular translation is unlikely in the apoplast, EVacs could potentially facilitate the cross-kingdom transfer of pre-assembled mRNA-ribosome complexes, allowing the plant to modulate the proteome of interacting pathogens or symbionts (*28*).

### EVacs originate from inward folding of the tonoplast, sequestrating cytoplasm

The stability of the GFP fluorescence in EVacs tagged with γ-TIP-GFP indicates that the GFP tag is not exposed to the acidic environment in either the vacuole or apoplast. This suggests that EVacs are unlikely to form via outward budding of the tonoplast, which would orient the C-terminus toward the apoplast. Plant vacuoles frequently generate tonoplast-derived intravacuolar spherical bodies through a process of tonoplast invagination and folding (fig. S17). These structures, which include vacuolar bulbs and other intralumenal invaginations, sequester cytoplasmic components (*21, 29, 30*). Once formed, these compartments remain functionally distinct and do not freely exchange proteins with the surrounding tonoplast (*20*). While they are marked by γ-TIP and other tonoplast markers, including Vam3/SYP22 (*31–33*), the exclusion of some tonoplast markers, such as Rab75c, highlights a selective protein segregation during their biogenesis (*21*).

Our live-imaging data showed that these tonoplast-derived intravacuolar bodies can be found outside the vacuole (fig. S18A and movie S8). They can move from the vacuolar lumen to the cytoplasm (fig. S18B), or form directly in the cytosol when the tonoplast folds upon itself (fig. S18C). Our data indicate that these bodies are subsequently secreted to the extracellular space as EVacs. Although a ‘direct-discharge’ model involving localized fusion of the tonoplast and PM is possible, the predominantly tonoplast-derived composition of the EVac membrane suggests that cytoplasmic tonoplast-derived bodies bud into the apoplast, encapsulated in plasma membrane (See image 1 of Fig. 2). In support of this model, proteomic analysis of the P10 fraction revealed multiple plasma membrane markers such as proton pumps (AHA1 and AHA2) and plasma membrane aquaporins (PIPs) (Data S1), and cryo-EM images frequently revealed a double membrane structure (Fig. 3C).

Collectively, our findings reveal a novel secretory pathway employed by plants that is mediated by EVacs, which function as large secretory organelles to transport proteins, RNAs, chloroplasts, vesicles, and other cytoplasmic components into the apoplast (fig. S19). This mechanism provides a plausible explanation for the long-standing observation that many apoplastic proteins lack canonical signal peptides and therefore cannot be secreted by the classical ER–Golgi secretory route. This unconventional protein secretion pathway in plants has previously been proposed to be mediated, at least in part, by exocyst-positive organelles (EXPOs), which are defined as double-membraned vesicles 500-800 nm in diameter located in the cytoplasm (*34, 35*). Secretion is accomplished by fusion of the outer membrane of EXPOs with the plasma membrane to release a single-membraned vesicle. The mechanism by which EXPOs are formed has not been defined. EXPOs might represent an intermediary stage in the secretion of EVacs. Although it is not yet clear how EVacs escape the vacuole, we propose four potential mechanisms (fig. S19). The first would involve budding out of the vacuole to form a double-membraned cytosolic vesicle similar to EXPOs, which would then be secreted by fusion of the outer membrane with the plasma membrane, generating an extracellular vesicle with a single lipid bilayer. The second would be for these EXPO-like vesicles to be directly budded from the plasma membrane, generating vesicles with three lipid bilayers, with plasma membrane on the outside. A third possible mechanism would be direct release from the vacuole through a transient pore formed by localized fusion of the tonoplast and plasma membrane. A fourth option could involve a process called hemifusion, in which two lipid bilayers come together and generate a single new bilayer (*36*). Such hemifusion could allow translocation of the intravacuoloar bulb to the cytoplasm then transfer from the cytoplasm to the apoplast, generating a vesicle with a single lipid bilayer with the outer leaflet derived from the plasma membrane and the inner leaflet derived from the tonoplast. Based on our cryo-EM imaging of purified EVacs, we observed EVacs with single lipid bilayers as well as EVacs with two lipid bilayers, or at least sections with two lipid bilayers (fig. S12), suggesting that there may be multiple mechanisms enabling EVac release. Regardless of the mechanism of EVac secretion, our data establish EVacs as a central player in the secretion of macromolecules and organelles from plant cells. It will be of interest to assess whether other organisms employ similar mechanisms, in particular, humans, as extracellular RNA is abundant in both human cell cultures and human plasma (*37, 38*).

## Acknowledgments

We thank Laurence Drouard for providing seed of *Arabidopsis rns* mutant lines, Mads Nielsen for providing the *Arabidopsis* RFP-PEN1 transgenic line, and Leonor Boavida for the TET8-GFP line. We also thank the *Arabidopsis* Biological Resource Center at The Ohio State University for providing the γ-TIP-GFP and T-DNA insertion lines. We also thank the Indiana University Bloomington Electron Microscopy Center for access to a Thermo-Fisher Talos-Arctica cryo-electron microscope and the Indiana University Light Microscopy Imaging Center for access to Leica SP8 and Stellaris confocal microscopes. The expert assistance of Barry Stein and David Morgan with electron microscopy, and Andras Kun with confocal microscopy is also gratefully acknowledged. We also thank Jonathan Trinidad and the IU Biological Mass Spectrometry Center for providing mass-spectrometry analyses.

## Funding

National Institutes of Health grant R35GM157146 (RJV)

National Institutes of Health grant T32 GM131994 (CTV)

National Science Foundation grants IOS-1842685, IOS-2141969 and IOS-2243531 (RWI)

National Science Foundation grant IOS-1842698 (BCM, PB)

National Science Foundation grant IOS-2243534 (PB)

Novo Nordisk Foundation (RWI)

## Author contributions

Conceptualization: MLB, MSR, MHSR, AYL, PB, RWI

Methodology: MLB, MSR, MHSR, AYL, GGG, SJCV, CTV, AF

Investigation: MLB, MSR, MHSR, GGG, CTV, AF

Visualization: MLB, MSR, MHSR

Funding acquisition: RJV, BCM, PB, RWI

Project administration: RWI

Supervision: RJV, BCM, PB, RWI

Writing – original draft: MLB

Writing – review & editing: MLB, MSR, MHSR, AYL, PB, AF, RJV, BCM, RWI

## Competing interests

RWI serves on the scientific advisory board for Terrana Biosciences, which is pursuing RNA-based solutions to promote plant health. All other authors declare that they have no competing interests.

## Data, code, and materials availability

All data are available in the main text or the supplementary materials.

## Supplementary Materials

### Materials and Methods

#### Plant materials and growth conditions

Seeds of *Arabidopsis thaliana* ecotypes Columbia-0 (Col-0) and Wassilewskija (Ws), wild type and related genotypes were sown on Sungro Propagation Mix, treated with fungicide (RootShield Plus), and stratified at 4 °C for 72 hours. Plants were grown in a controlled chamber at 24 °C under short-day conditions (9-h light/15-h dark) with a light intensity of 120 µmol m⁻² s⁻¹. Seedlings were transplanted after 10 days into individual 36-cell tray inserts containing Sungro Professional Growing Mix and fertilized periodically with Miracle-Gro All Purpose Plant Food (24-8-16) at a concentration of 250 mg/L.

The *atg5*-1 (*39*), *sid2-2* (*40*), and *sid2/atg5* (*16*) mutants were previously described. The *rns1* (FLAG_566A08), *rns3* (FLAG_164A04), and the double mutant *rns1rns3* (*10*) in the Ws background were kindly provided by Dr. Laurent Drouard (IBMP, Strasbourg). The 35S::RFP-PEN1 line (*41*) was provided by Mads Nielsen (Copenhagen University) and the native promoter TET8-GFP line (*42*) was provided by Leonor Boavida (Purdue University). The following lines in the Col-0 background were obtained from the *Arabidopsis* Biological Resource Center (ABRC): *rns2-2* (SALK_069588), γ-TIP-GFP (CS16257), γ-TIP-CFP (CS16256), and γ-TIP-YFP (CS16258).

The dual-labeled line RFP-PEN1 x γ-TIP-GFP was generated by genetic crossing. The γ-TIP-GFP TET8-mCherry reporter line was generated by Agrobacterium-mediated transformation of the γ-TIP-GFP reporter line using a pEG100 vector containing 35S::TET8-mCherry via the floral dip method (*43*). Transgenic seedlings were selected on MS plates supplemented with 50 mg/L Glufosinate-ammonium (BASTA).

#### Isolation of Apoplstic Wash Fluid (AWF)

AWF was extracted from 6 week-old plants following the stepwise procedure described by (*44*) with specific modifications regarding buffer composition. Fresh rosettes were detached and rinsed several times in distilled water to remove soil particles, and vacuum-infiltrated twice for 20 s with VIB. Infiltrated rosettes were carefully blotted dry and placed in 60-mL syringes nested within 250-mL centrifuge bottles and centrifuged at 600 *g* for 30 min at 4 °C. To investigate the effect of pH on RNA stability, the MES (pH 6.0) in the VIB was replaced with either 20 mM HEPES (pH 8.2) or a 15 mM Carbonate-bicarbonate buffer pH 9.2 (18.2 mM NaHCO_3_, 1.8 mM Na_2_CO_3_). AWF was then filtered through a 0.2 µm syringe filter or alternatively unfiltered for EVac isolation.

#### Isolation of EVacs from AWF

Extracellular vacuole-derived bodies (EVacs) were isolated from unfiltered AWF. AWF was aliquoted into 2-mL microcentrifuge tubes (1.5 mL per tube) and subjected to centrifugation at 4 °C for 35 min. EVac-enriched fractions were obtained at 2,000 *g* (P2), 10,000 *g* (P10), or 15,000 *g* (P15). In specific experiments, a differential centrifugation was performed where the supernatant from the 2,000 *g* spin was further centrifuged at 15,000 *g* to recover the remaining EVac population (P15-P2). The absence of EVacs in the resulting supernatants S10 and S15 confirmed that 35 min of centrifugation at 10,000–15,000 *g* is sufficient to recover essentially all EVacs.

Following centrifugation, the supernatant was decanted and the tube walls were carefully dried with a Kimwipe. Pellets were resuspended gently in 100 µL of either VIB (pH 6.0) or 20 mM HEPES (pH 8.2; supplemented with 2 mM CaCl₂ and 10 mM NaCl) using wide-bore pipette tips (manually cut) to prevent the mechanical rupture of the fragile EVacs. For RNA analysis, resuspended pellets were immediately processed with TRIzol reagent. For immunoblot analysis, 30 µL of the EVac suspension was immediately mixed with 10 µL of 4x SDS loading buffer (250 mM Tris-HCl pH 6.8, 8% SDS, 40% glycerol, 20% 2-mercaptoethanol, and 0.004% bromophenol blue), denatured at 95°C for 5 min or 65°C for 15 min, and stored at –80°C until further use. For confocal imaging, 20–30 µL of the EVac suspension was diluted with 40 µL of the corresponding buffer and loaded into 18-well chambered coverslips (1.5H ibidi Glass Bottom). To facilitate EVac adherence, coverslips were manually coated with poly-D-lysine and used on the day of preparation.

#### Visualization of RNA inside EVacs

EVacs were isolated from the γ-TIP-CFP transgenic line as described above. The resulting pellets were gently resuspended in 80 µL of 1X PBS (pH 7.4) or 20 mM HEPES pH 8.0 and incubated with 500 nM SYTO RNASelect (Invitrogen, S32703) for 20 min at room temperature in the dark. For imaging, 30-40 µL of the labeled suspension was loaded onto poly-D-lysine - coated 18-well chambered coverslips (ibidi). Confocal microscopy was performed using a Leica Stellaris 8 system equipped with a White Light Laser (WLL) and HyD detectors. Images were acquired on a Leica Stellaris 8 system using an HC PL APO 63x/1.20 water immersion objective. CFP fluorescence was detected using an excitation of 440 nm with an emission range of 445–497 nm. SYTO RNASelect was excited at 490 nm with emission collected at 500–548 nm. Images were processed in Fiji in the same way as the Giant Unilamellar Vesicles (GUVs) used as positive and negative controls.

#### Preparation and imaging of Giant Unilamellar Vesicles (GUVs)

Giant unilamellar vesicles (GUVs) were generated via electroformation as previously described (*45*), with minor modifications. A lipid mixture consisting of cholesterol:POPC:POPE:POPS (11:55:27:7 molar ratio) was prepared in chloroform. A 30-µL aliquot of the lipid solution was deposited as a thin film onto the conductive side of unpolished ITO-coated glass slides (Structure Probe, Inc.) and dried under vacuum (30 inHg) at 50 °C for 30 min. The lipid film was rehydrated within a silicone isolator (Grace Bio-labs) using 500 µL of either 300 mM sucrose (negative control GUVs) or 2.5 µg/µL of total leaf *Arabidopsis* RNA in 300 mM sucrose (RNA-containing GUVs). Electroformation was conducted at room temperature for 1.5 h using a 3 V peak-to-peak sine wave at 10 Hz, followed by 10 min at 2 Hz to facilitate GUV detachment from the slides. Harvested GUVs were diluted 1:1 in 300 mM sucrose and imaged on the same day.

For confocal microscopy, 5 µL of the GUV suspension was added to an 8-well chambered coverslips (ibidi) containing 265 µL of 1x PBS pH 7.4 and incubated with 500 nM SYTO RNA Select for 20 min. GUV membranes were counterstained with CellMask Deep Red (5 µg/mL) for 5 min prior to imaging. Images were acquired on a Leica Stellaris 8 system using an HC PL APO 63x/1.20 water immersion objective. SYTO RNASelect was excited at 490 nm (emission 500– 548 nm), and CellMask Deep Red was excited at 650 nm (emission 670–697 nm). Images were processed in Fiji, with negative controls processed in the same way as positive controls.

#### Ribonuclease protection assay

To confirm the encapsulation of RNA within EVacs, an RNase protection assay was performed under osmoprotective conditions. EVacs (P10) were isolated using VIB supplemented with 0.4 M sorbitol to maintain membrane integrity and minimize mechanical rupture during incubation. Samples were maintained on ice throughout the procedure unless otherwise specified.

The P10 pellet was resuspended in 150 µL of VIB + 0.4 M sorbitol. Resuspended EVacs were split into 50 µL aliquots and the following treatments were performed: (i) Detergent + RNase A, where membranes were solubilized with 1.0% (v/v) Triton X-100 for 40 min at room temperature prior to enzyme addition; (ii) RNase A alone, to digest non-encapsulated RNA; and (iii) Mock treatment, where EVacs were incubated with buffer only.

For RNase treatments, samples were incubated with 0.1 µg/mL RNase A (diluted in 10 mM Tris-HCl pH 7.5 and 15 mM NaCl) for 45 min on ice. The detergent-treated samples served as a positive control for digestion, to validate that the RNase A concentration was sufficient to degrade RNA once the lipid bilayer was disrupted. All reactions were stopped by the addition of 1 mL of TRIzol reagent, followed by immediate RNA extraction.

#### Ribonuclease activity assays

To evaluate the ribonuclease activity of extracellular fluids, 500 ng of total leaf RNA (isolated using TRIzol) was incubated with 50 µL of AWF. Samples were incubated at room temperature with agitation for 0, 10, 30, and 60 min. Reactions were stopped by adding 1 mL of TRIzol reagent, followed by RNA extraction as described below. The endogenous RNA content within the AWF volume used for these assays was determined to be negligible compared to the 500 ng of total cell lysate RNA.

#### Isolation of intact vacuoles from *Arabidopsis* protoplasts

Intact vacuoles were isolated following a modified version of the osmotic and thermal disruption method (*46*). Protoplasts were first generated using the “tape-sandwich” method (*47*) to minimize mechanical stress. Briefly, the abaxial epidermal layer of 5- to 6-week-old leaves was removed using adhesive tape, and the exposed mesophyll was incubated in a filter-sterilized enzyme solution (1.5% [w/v] Cellulase R-10, 0.4% [w/v] Macerozyme R-10, 0.4 M mannitol, 20 mM KCl, 10 mM CaCl₂, 0.1% BSA, 0.035% β-mercaptoethanol, and 20 mM MES, pH 5.7).

After a 3 h incubation at room temperature in the dark, protoplasts were carefully transferred to a 50-mL centrifuge tube and collected by centrifugation (100 *g*, 5 min, RT). The pellet was washed with wash buffer (0.4 M mannitol and 10 mM MES pH 5.7), followed by centrifugation at 100 *g* for 3 min.

To release the vacuoles, the purified protoplast pellet was resuspended in a pre-heated (37°C) lysis buffer (0.2 M mannitol, 10% [w/v] Ficoll 400, and 10 mM EDTA in 5 mM Na₂HPO₄/NaH₂PO₄ buffer, pH 8.0). The resulting lysate was subjected to a discontinuous Ficoll density gradient: 7 mL of the lysate was overlaid with 3 mL of 4% (w/v) Ficoll (prepared in Vacuole Buffer: 0.45 M mannitol, 2 mM EDTA, and 5 mM Na₂HPO₄/NaH₂PO₄, pH 7.5) and a final top layer of 1 mL of ice-cold Vacuole Buffer. The gradient was centrifuged in a SW 41 Ti swinging-bucket rotor at 71,000 *g* for 50 min at 10°C. Intact vacuoles were recovered from the interface between the Vacuole Buffer and the 4% Ficoll layers. To preserve their structural integrity, all handling was performed using manually cut wide-bore pipette tips. Isolated vacuoles were either used immediately for confocal imaging or aliquoted and stored at –80 °C for subsequent analysis.

#### RNA extraction and purification

RNA was isolated from various plant fractions using TRIzol or TRIzol LS Reagents (Thermo Fisher Scientific) according to the sample volume and type. For total cell lysate RNA, approximately 100 mg of leaf tissue was frozen in liquid nitrogen, ground to a fine powder, and homogenized in 1 mL of TRIzol. For isolated vacuoles and extracellular fractions (AWF, EVac fractions and supernatants) with volumes ≤100 µL, 1 mL of TRIzol was used directly. For liquid samples with volumes between 100 µL and 250 µL, TRIzol LS was employed at a 3:1 (v/v) reagent-to-sample ratio. For liquid fractions exceeding 250 µL (e.g., AWF), RNA was first concentrated by precipitation with 0.1 volumes of 3 M sodium acetate (pH 5.2) and one volume of cold isopropanol at –20 °C (1 h to overnight), followed by centrifugation at 15,000 *g* for 30 min at 4 °C. The resulting pellets were resuspended in 100 µL of ultrapure water and subsequently processed with 1 mL of TRIzol.

Phase separation was achieved by adding 200 µL of chloroform per 1 mL of TRIzol/ 0.75 mL TRIzol LS, followed by vortexing and centrifugation at 13,000 *g* for 15 min at 4 °C. To exclude potential Ficoll carryover in purified vacuole samples, 0.5 volumes of 7.5 M ammonium acetate were added to the recovered aqueous phase prior to isopropanol precipitation. For all samples, the aqueous phase was supplemented with 10 µg of RNase-free glycogen and one volume of cold isopropanol and incubated 1 h at –20°C. RNA was pelleted at 15,000 *g* for 20 min at 4°C, washed twice with cold 70% ethanol, and resuspended in ultrapure water.

When necessary, RNA samples were further purified to remove residual phenol or guanidine contamination by serial precipitation. Briefly, 1 µg of glycogen and 0.5 volumes of 7.5 M ammonium acetate were added to the RNA, followed by 2.5 volumes of 100% ethanol. Mixtures were incubated at –80°C for 30 min, centrifuged at 13,000 *g* for 20 min at 4°C, and the final pellets were washed with cold 70% ethanol.

#### RNA quantification

RNA concentrations were determined using a combination of spectrophotometry and fluorescence-based assays. For total cell lysate RNA, initial concentrations were estimated using a NanoDrop spectrophotometer (Thermo Fisher Scientific). However, as previously reported (*6*), extracellular samples contain non-nucleic acid contaminants that lead to the overestimation of RNA levels by absorbance-based methods. ExRNA fractions were quantified using a SYBR Gold fluorescence assay in a microplate reader, following the protocol described (*44*). Briefly, 1 µL of RNA was incubated with 150 µL of 0.5X SYBR Gold Nucleic Acid Gel Stain (Invitrogen) in 200 mM Tris pH 8.0 for 5-10 minutes, and fluorescence was measured using a Synergy H1 microplate reader (excitation 496 nm, emission 540 nm). Concentrations were calculated against a standard curve generated with known amounts of total cell lysate *Arabidopsis* RNA. For specific experiments, relative quantifications were performed by densitometric analysis of 12% or 15% Urea-PAGE gels stained with 1X SYBR Gold, using ImageJ software to calculate the pixel intensity as previously described (*6, 44*).

#### Denaturing Polyacrylamide Gel Electrophoresis (PAGE) of RNAs

Electrophoresis was performed using either 15% precast TBE-Urea gels (Novex, Invitrogen) or hand-cast 12% polyacrylamide gels containing 7 M urea in 1X Tris-Boric Acid-EDTA (TBE, pH 8.4). Hand-cast gels were prepared using 40% Acrylamide/Bis Solution (37.5:1; Bio-Rad) and cast in either Novex empty cassettes or the Mini-PROTEAN system (Bio-Rad).

For comparisons among EVac pellets (P2, P10, and P15), loading was normalized according to equal starting AWF volume. For comparisons between EVac pellets and their corresponding supernatants or between EVac pellets and vacuolar fractions, supernatant and vacuole lanes were loaded with equal amounts of RNA relative to the pellets. RNA samples were mixed 1:1 with 2X denaturing loading buffer (95% formamide, 10 mM EDTA, 0.02% SDS, 0.02% bromophenol blue, and 0.01% xylene cyanol), denatured at 65 °C for 10 min, and resolved in 0.5X TBE running buffer at room temperature. For size estimation, a 1:1 mixture of Low Range ssRNA Ladder (New England Biolabs) and 14–30 nt ssRNA Ladder Marker (Takara) was used. Gels were stained with 1X SYBR Gold Nucleic Acid Gel Stain (Invitrogen) in 0.5X TBE for 10 min, washed twice with distilled water or 0.5X TBE, and imaged using a Bio-Rad ChemiDoc system.

#### DIG-labeled RNA gel-blots

For detection of specific tRNA and rRNA species, RNAs resolved by PAGE were transferred onto positively charged nylon membranes (Hybond-N+, Cytiva) using a semi-dry Trans-Blot Transfer System (Bio-Rad) at a constant 20 V for 45 min in 0.5X TBE. Membranes were UV cross-linked twice at 120,000 µJ/cm² for 30 s (UVC-508, Ultra-Lum) and prehybridized for at least 40 min at 42 °C in DIG Easy Hyb solution (Roche) supplemented with 0.1 mg/mL Poly(A).

Hybridization was performed overnight at 42°C with 2.5 pmol/mL digoxigenin (DIG)-labeled DNA probes in DIG Easy Hyb solution containing Poly(A). DNA oligonucleotides (IDT) were 3’-end labeled using the DIG Oligonucleotide Tailing Kit (2nd Generation, Roche) according to the manufacturer’s instructions. Following hybridization, membranes were washed twice in low-stringency wash buffer (2X SSC, 0.1% SDS) at room temperature for 5 min, and twice in high-stringency wash buffer (1X SSC, 0.1% SDS) at 42°C for 15 min. Membranes were blocked for 40 min in 1X Blocking Solution (Roche) and incubated for 30 min with an alkaline phosphatase-conjugated anti-DIG antibody (Roche). Chemiluminescent signals were developed using CDP-Star (Roche) and captured with a Bio-Rad ChemiDoc imaging system. Oligonucleotide sequences used as probes are listed in Table S1.

#### Immunoblots

Leaf tissues (150 mg) were flash-frozen in liquid nitrogen and ground to a fine powder and proteins were extracted using cold extraction buffer (150 mM NaCl, 50 mM Tris-HCl pH 7.5, 0.1% [v/v] NP-40, 1% DPDS and 1% [v/v] plant protease inhibitor cocktail). The lysate was then centrifuged at 15,000 *g* for 10 min at 4°C and the supernatant used for subsequent analysis.

Protein concentrations were quantified using the Pierce 660 nm reagent (Thermo Fisher Scientific) against a bovine serum albumin (BSA) standard curve. AWF, purified vacuoles, and EVac fractions, as well as their respective supernatants, were used directly for analysis. For comparisons among EVac pellets (P2, P10, and P15), loading was normalized according to equal starting AWF volumes. For comparisons between EVac pellets and their corresponding supernatants, the supernatant was loaded based on equal protein amounts relative to the pellets. When specific compartment fractions (AWF, EVac pellet, supernatant or vacuole) were compared to cell lysate (CL), a higher protein amount was loaded for CL to enable detection of compartment-specific markers that are diluted in the total cellular context. Protein samples were mixed in a 3:1 ratio with 4X loading buffer (250 mM Tris-HCl pH 6.8, 8% [w/v] SDS, 40% [v/v] glycerol, 0.004% bromophenol blue, and 20% [v/v] 2-mercaptoethanol). Samples were denatured at 95 °C for 5 min or 65 °C for 15 min and resolved on 4–20% TGX stain-free pre-cast gels (Bio-Rad). Electrophoresis was conducted at 120-150 V for about 1 h. Stain-free images were captured using a Bio-Rad ChemiDoc imaging system. Proteins were transferred to 0.45 µm nitrocellulose membranes (Amersham Protran) using a conventional tank (wet) transfer system (Bio-Rad) in transfer buffer (25 mM Tris, 200 mM glycine, 20% [v/v] methanol) at 300 mA for 1 h. Proteins transferred to the membrane were visualized by applying Ponceau Stain (0.1% w/v Ponceau S Stain, 5% v/v acetic acid) for 10 min and rinsing in distilled water.

Membranes were blocked in 5% (w/v) skim milk (Difco, BD) in TBST (100 mM Tris, 150 mM NaCl, 0.1% Tween-20, pH 7.5) for 1.5 h at room temperature. Primary antibody incubations were performed overnight at 4 °C using the following dilutions: anti-PEN1 (1:1,000; (*48*)), anti-VHA-E1 (1:1,000; Agrisera, AS07 213), anti-GFP (1:2,000; Sigma, 11814460001), anti-Aleurain (1:1,000; Agrisera, AS20 4406), anti-RNS3 (1:1,000; Agrisera, AS22 4859), anti-RFP (1:2,000, ChromoTek, 6g6) and anti-Rps6 (1:2,000; (*49*)). Membranes were washed with TBST and incubated with HRP-conjugated secondary antibodies (goat anti-rabbit or anti-mouse at 1:8,000) for 1.5 h at room temperature. Signals were developed using ProtoGlow ECL substrate (National Diagnostics) and captured with a ChemiDoc system.

#### Mass Spectrometry (MS)

For protein digestion, two independent biological replicates of pelleted EVac fractions (P10), isolated and analyzed on separate days, were denatured in 8 M urea in 100 mM ammonium bicarbonate. Samples were incubated for 45 min at 57°C with 10 mM Tris(2-carboxyethyl)phosphine hydrochloride to reduce cysteine residue side chains, followed by alkylation with 20 mM iodoacetamide for 1 h in the dark at 21°C. The urea concentration was diluted to 1 M using 100 mM ammonium bicarbonate. A total of 0.4 μg of trypsin (Promega) was added, and samples were digested for 14 h at 37°C.

The resulting peptide solution was desalted using ZipTip pipette tips (EMD Millipore), dried down, and resuspended in 0.1% formic acid. Peptides were analyzed by LC-MS/MS on an Orbitrap Fusion Lumos mass spectrometer equipped with an Easy NanoLC 1200 (Thermo Fisher Scientific) at the Laboratory for Biological Mass Spectrometry (Indiana University Bloomington). Buffer A consisted of 0.1% formic acid in water, and Buffer B consisted of 0.1% formic acid in 80% acetonitrile. Peptides were separated on a 90-min gradient from 0% B to 35% B. Precursor ions were measured in the Orbitrap with a resolution of 60,000 and fragmented by higher-energy collisional dissociation (HCD) at a relative collision energy of 32%. Fragment ions were analyzed in the Orbitrap at a resolution of 15,000.

Data analysis was performed using Proteome Discoverer (v2.5, Thermo Fisher Scientific). MS/MS spectra were searched against the *Arabidopsis thaliana* TAIR10 proteome database, downloaded 02/2015. Trypsin was specified as the protease, allowing up to two missed cleavages. Carbamidomethylation of cysteine was set as a fixed modification, while methionine oxidation and protein N-terminal acetylation were included as variable modifications. A precursor mass tolerance of 10 ppm and a fragment ion tolerance of 0.04 Da were applied. Peptide and protein identities were validated using the Percolator node. All identified proteins meeting high and medium False Discovery Rate (FDR) confidence thresholds were retained for subsequent analysis.

#### Live Confocal Fluorescence Microscopy

Confocal fluorescence microscopy was performed using Leica Stellaris 8 and Leica SP8 systems equipped with HC PL APO 40x/1.10 and 63x/1.20 water immersion objectives. Imaging was conducted on fresh leaves from either 2-week-old seedlings or 6-week-old adult rosettes. To enhance the visualization and staining of the mesophyll layer, in some instances the abaxial epidermis of freshly detached leaves was carefully removed using the “tape-sandwich” method (*47*) prior to imaging. For plasma membrane labeling, leaves were incubated with 5 µg/mL CellMask Orange (Thermo Fisher) in VIB for 10 min at room temperature, followed by three washes in buffer. To eliminate apoplastic air and minimize light reflection, leaves were vacuum-infiltrated for 10-15 sec in distilled water or VIB. Infiltrated leaves were mounted on coverslips and visualized within 2 h.

Freshly isolated EVac preparations were visualized by mounting 30-50 µL of the fraction directly onto poly-D-lysine-coated 8-well μ-Slide chambered coverslips (ibidi). For EVacs isolated from Col-0 plants, membranes were stained with 0.5 µg/mL CellMask Deep Red (Thermo Fisher) for 10 min prior to imaging.

Fluorescence was detected using the following excitation/emission (Ex/Em) parameters: CFP (Ex:440; Em:445–497 nm), GFP (Ex: 488 nm; Em: 498–550 nm), YFP (Ex: 514 nm; Em: 524–560 nm), RFP (Ex: 543 nm; Em: 566–630 nm), mCherry (Ex: 586 nm; Em: 597–625 nm), CellMask Orange (Ex: 553 nm; Em: 561–601 nm) and chlorophyll autofluorescence (Ex: 488 nm; Em: 655–730 nm. Unless otherwise specified, images represent single optical planes. Raw images were processed and analyzed using the FIJI/ImageJ software package.

#### Cryo-EM and 2D class averaging

Cryo-EM grid preparation was performed using two independent biological replicates prepared and imaged on separate days at the Electron Microscopy Center (Indiana University Bloomington). Lacey carbon-coated nickel grids (LC200-Ni; Electron Microscopy Sciences) were glow-discharged to render the carbon film hydrophilic prior to sample application. Cryo-EM grids were prepared by applying 2-3 µL of the isolated EVac fraction (P10) inside the environmental chamber of a Vitrobot plunge-freezing device (Vitrobot Mark IV, Thermo Fisher Scientific) maintained at 22°C and 100% relative humidity. Vitrification was achieved utilizing a single blot with a blotting time of 4.0 or 6.0 s, a blotting force of 4, 6 or 8, a wait time of 0 s, and a drain time of 0 s. Samples were immediately vitrified by plunge-freezing into liquid ethane cooled by liquid nitrogen. After grid screening, data collection was performed using a Talos Arctica cryo-transmission electron microscope at 200 keV (Thermo Fisher Scientific) equipped with a k3 camera (Gatan) and BioContinum energy filter (Gatan). Micrographs were obtained at nominal magnifications of 31k, 39k, 63k and 79k, exposure time of 3s, 30 frames per movie, total electron dose of 54.2 e−/ Å2 and −2 μm defocus. The 30 frames from each movie were averaged to generate a single micrograph, which was used for figure preparation and subsequent analysis. The pixel sizes were 2.803 Å, 2.213 Å, 1.372 Å, and 1.09 Å, respectively.

Seven micrographs containing non-vesicular particles were selected and imported to RELION 5.0.1 (*50*). The four nominal magnifications were assigned to four corresponding optics groups. Contrast Transfer Function (CTF) estimation was done by CtfFind-4.1.14 (*51*). A total of 573 particles were manually picked and extracted at final pixel sizes of 2.803 Å/pixel for 31k, 2.213 Å/pixel for 39k, 2.744 Å/pixel for 63k, and 2.18 Å/pixel for 79k. The particles were then subjected to three successive rounds of 2D classification with a 300 Å diameter circular mask. The numbers of classes (K) were 10, 10, and 5 for the three rounds, respectively. Of the final five classes, one contained 82 particles and represented the large-particle population (∼300 Å diameter), whereas another contained 85 particles representing the small particle population (∼150 Å diameter).

#### 3′ poly(A) RNA sequencing

RNA from whole leaf and EVac preparations (P10) was isolated as described above. The isolated RNA was shipped on dry ice to Plasmidsaurus (Louisville, KY, USA). Libraries were prepared using a stranded 3′ poly(A) RNA-sequencing workflow in which polyadenylated transcripts were captured by oligo(dT)-primed reverse transcription with unique molecular identifiers (UMIs), followed by second-strand cDNA synthesis, tagmentation, indexing, and PCR amplification. Libraries were sequenced on an Illumina platform to generate 3′ end-counting reads (https://plasmidsaurus.com/technical-documentation/rna).

#### Computational analysis

The RNAseq reads were subjected to the following quality control steps to obtain high quality (HQ) reads for further downstream analysis. Ribosomal RNA reads were removed using SortMeRNA against the SILVA database version 138.2 (*52, 53*). The remaining reads were filtered using FastP v0.23.2 (*54*) by performing poly-X tail trimming, 3′ quality-based trimming with a minimum Phred quality score of 15 and discarding reads shorter than 50 bp after trimming. The quality reports of raw and HQ reads were generated using FastQC (*55*).

HQ reads from cell lysate and EVac groups were aligned independently to the *Arabidopsis thaliana* genome (GCA_978657495.1) (https://www.ncbi.nlm.nih.gov/bioproject/?term=PRJEB100887) using STAR v 2.7.11b (*56*) with removal of non-canonical splice junctions. Gene-level read counts were generated using featureCounts v 2.0.6 with the TAIR12 genome annotation (*57*). Genes with ≥10 read counts in at least two biological replicates per condition were considered expressed. Overlap between conditions was visualized using R. Raw read counts were normalized to reads per million (RPM) using the total number of mapped reads for each sample. Principal component analysis (PCA) was performed on log_2_(RPM + 1) values using the prcomp function in R. Further, genes detected in at least four of six samples were retained for analysis. Genes were classified as cell lysate– enriched, EVac-enriched, Shared (highly expressed in both), or low abundance based on mean RPM values and fold-change threshold. Cumulative RPM values for each category were visualized as stacked bar plots in R. Genes expressed in at least four of six samples were classified as cell lysate–enriched (mean RPM ≥10; FC ≤0.5), EVac-enriched (mean RPM ≥10; FC ≥2), or Shared (highly expressed in both groups) (mean RPM ≥10 in both groups; 0.5 < FC < 2). The Genes with RPM ≤10 were classified as the low category. Heatmaps were generated in R using the pheatmap package from log_2_(RPM + 1) values of the 20 most highly expressed shared genes and the 20 most EVac-enriched genes, with hierarchical clustering of genes only.

Visualization of the data was done using R.

---

**Fig. S1.**
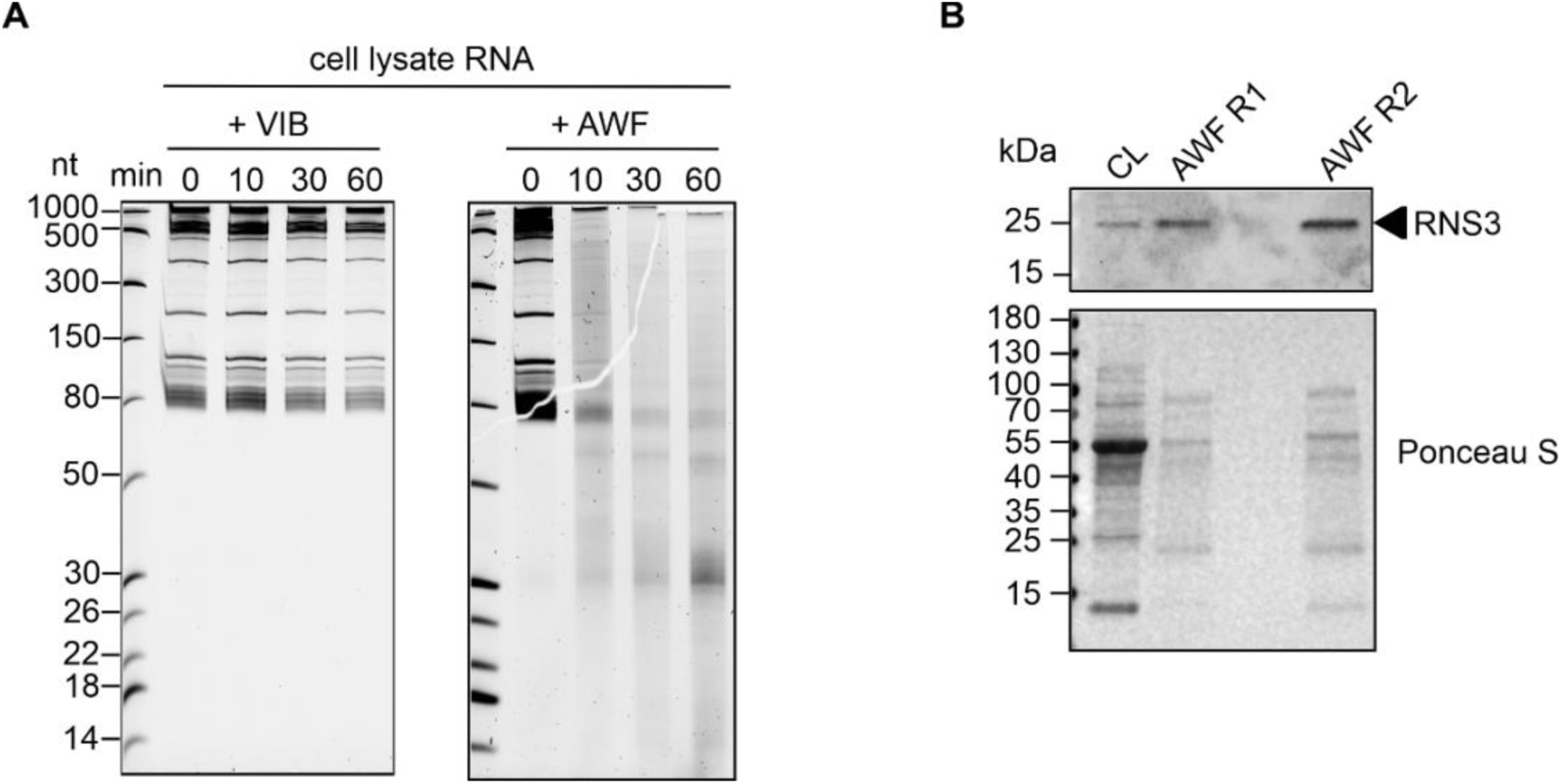
Apoplastic RNases rapidly degrade total cellular RNA. **(A)** Total cell lysate RNA was incubated at RT for the indicated times with Vesicle Isolation Buffer (VIB) or AWF. Reactions were stopped with TRIzol prior to RNA extraction and resolved by denaturing PAGE. RNA ladder sizes are indicated in nucleotides (nt). **(B)** Immunoblot showing RNS3 enrichment in AWF relative to total cell lysate (CL) protein. Loading is described in Materials and Methods.

**Fig. S2.**
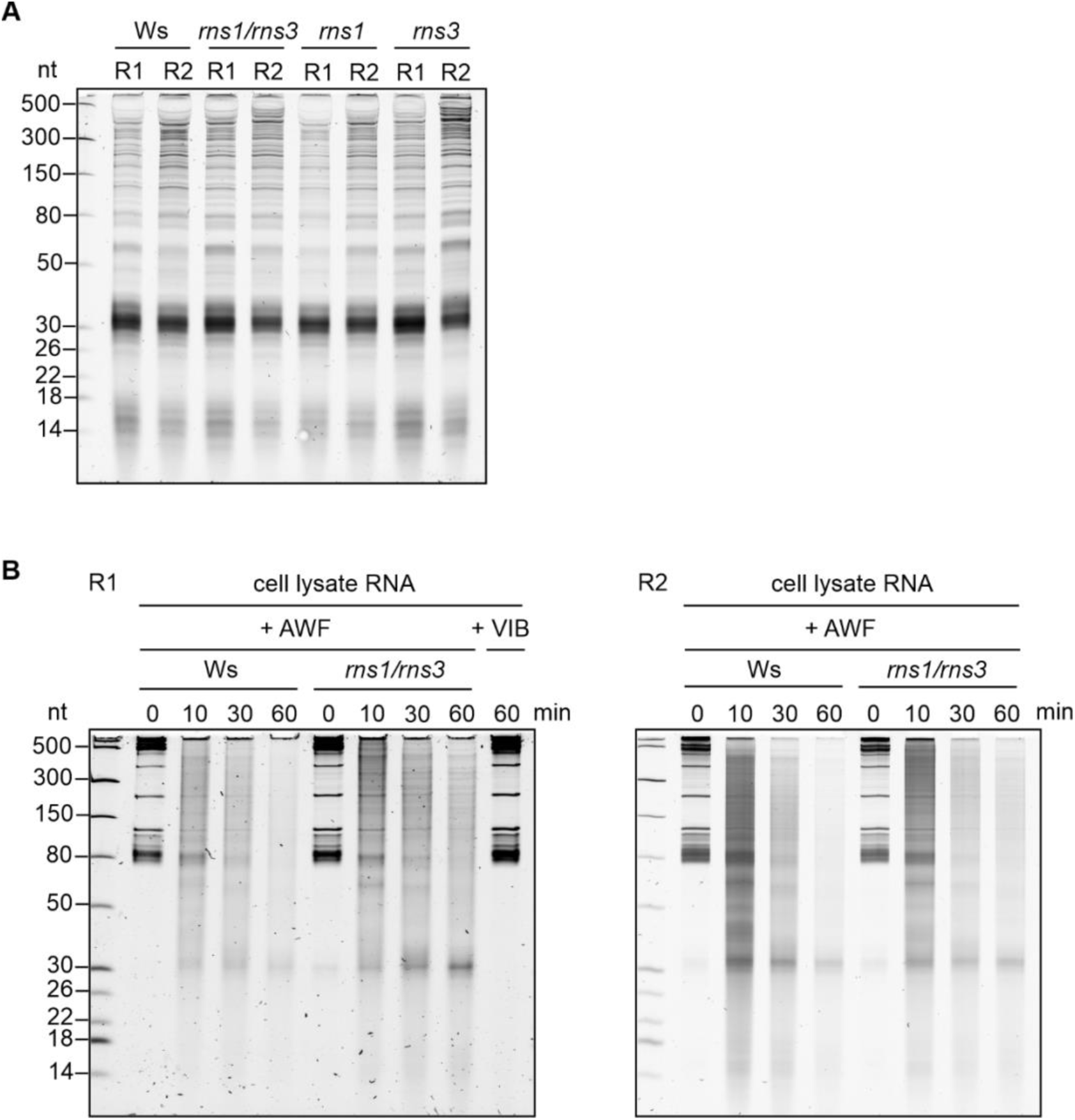
ExRNA processing is mediated by multiple redundant RNases. **(A)** The lack of RNS1 and RNS3 does not affect the apoplastic RNA pattern. RNA was isolated from equal volumes of AWF from the indicated genotypes using TRIzol and resolved by denaturing PAGE. Two biological replicates (R1 and R2) are shown. **(B)** The lack of RNS1 and RNS3 has a small impact on the ability of AWF to degrade naked cell lysate RNA. Total cellular RNA was incubated at RT with VIB or AWF from the indicated *Arabidopsis* genotypes for the indicated times prior to TRIzol extraction and denaturing PAGE. Two biological replicates (R1 and R2) are shown.

**Fig. S3.**
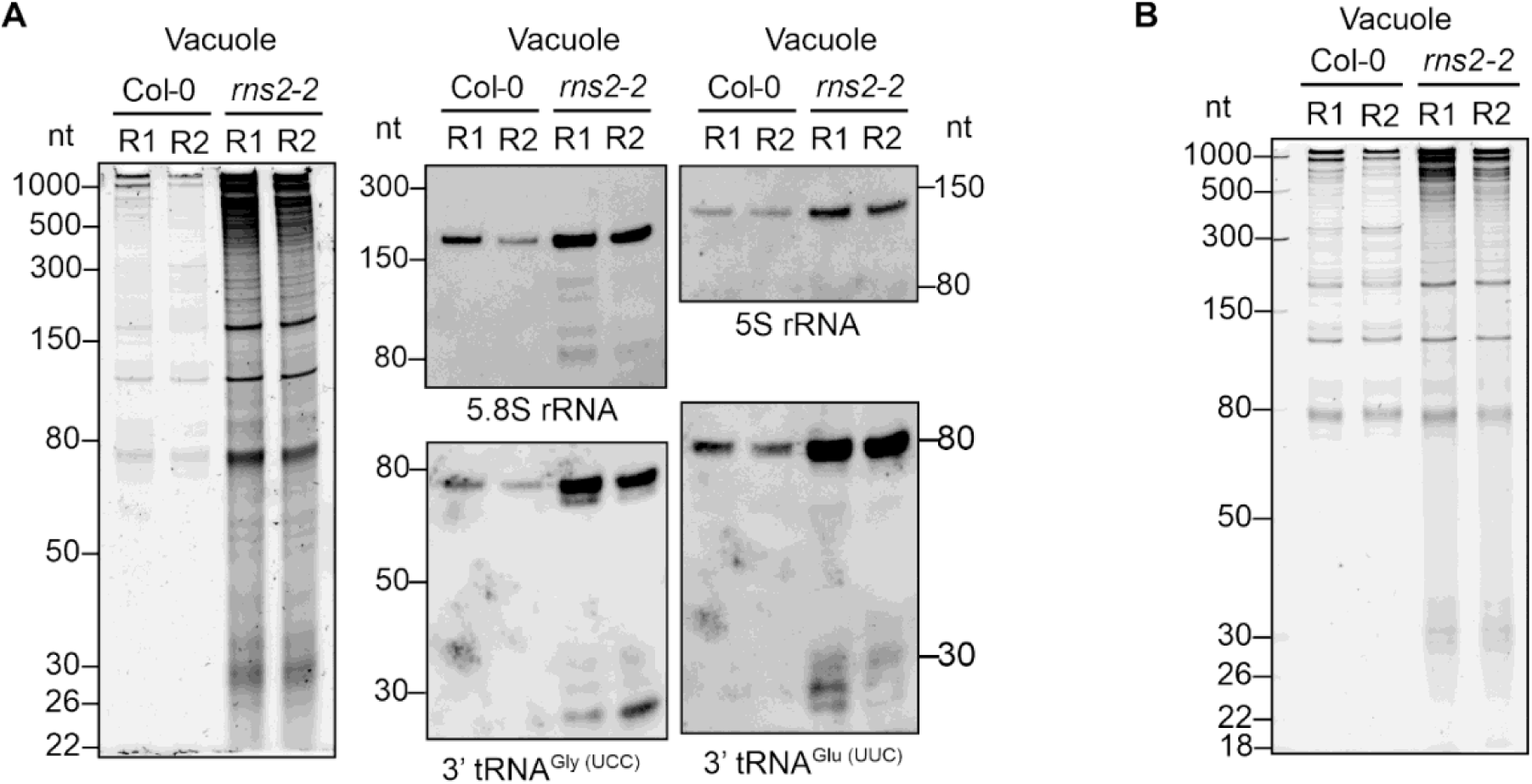
Vacuolar cargo contributes to the RNA composition of the apoplast. **(A)** RNA profile (left) and northern blot detection of extracellular tRNAs and rRNAs (right) in vacuoles isolated from Col-0 and *rns2-2* rosette leaves. Vacuoles were isolated from equal leaf fresh weights. Two biological replicates (R1-R2) are shown. **(B)** Denaturing PAGE of RNA isolated from vacuoles purified from Col-0 and *rns2-2* rosette leaves. Samples were loaded based on equal total RNA amounts to facilitate comparison between genotypes. Two biological replicates (R1-R2) are shown.

**Fig. S4.**
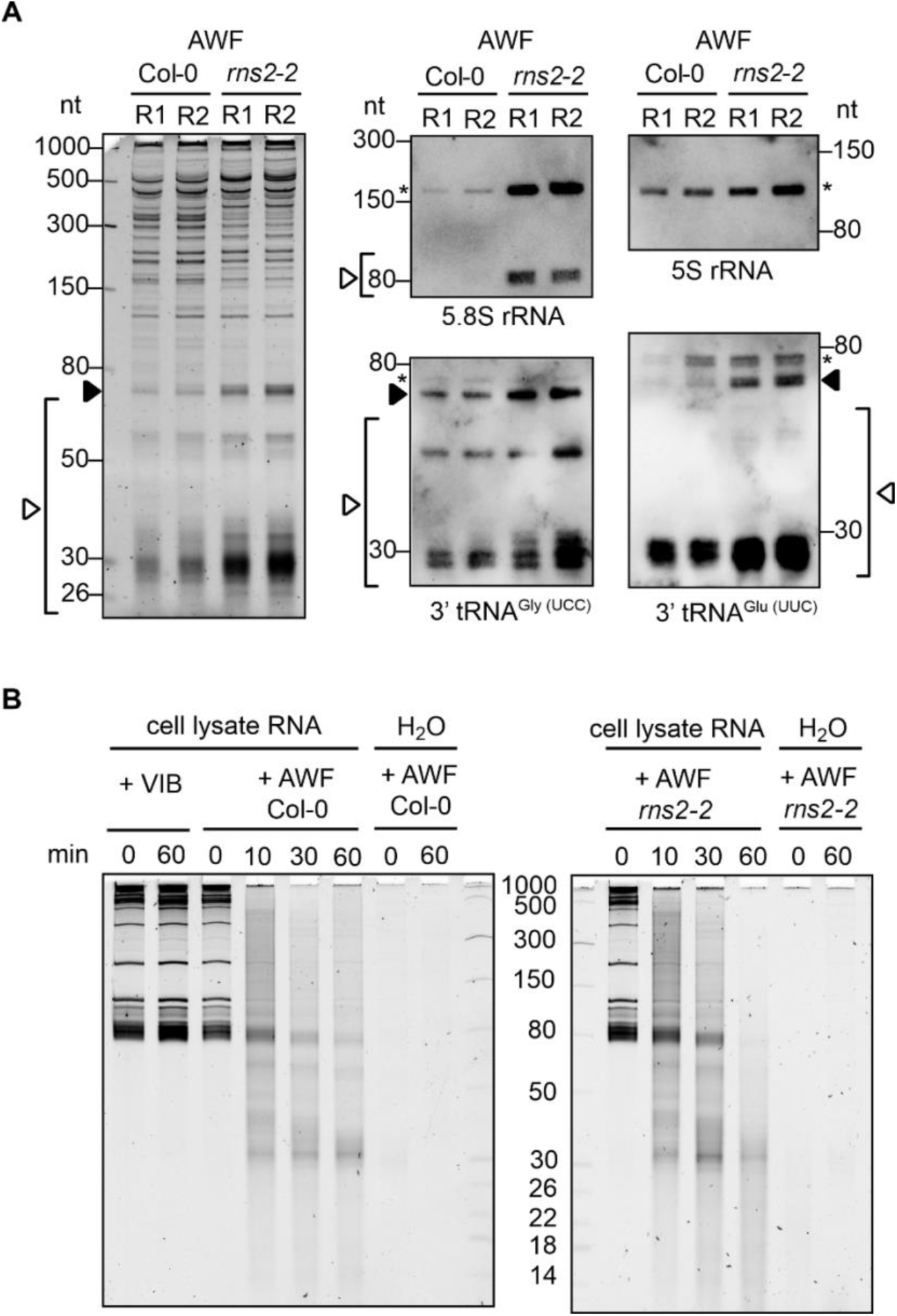
AWF-mediated RNA cleavage is maintained in the absence of RNS2. **(A)** RNA profile and RNA gel-blot detection of AWF isolated from Col-0 and *rns2-2* rosette leaves. RNA was isolated from an equal volume of AWF, corresponding to equal leaf fresh weight. Asterisks indicate full length rRNAs or tRNAs; black arrowheads indicate slightly truncated tRNAs and open arrowheads indicate tRNA and rRNA derived fragments. Two biological replicates (R1-R2) are shown. **(B)** RNA gel showing total cellular RNA incubated with AWF from wild-type Col-0 and *rns2-2* at RT for the indicated times. As a control, cell lysate RNA was incubated with VIB. RNA recovered from an equal volume of AWF incubated with dH_2_O is shown to indicate negligible endogenous AWF RNA.

**Fig. S5.**
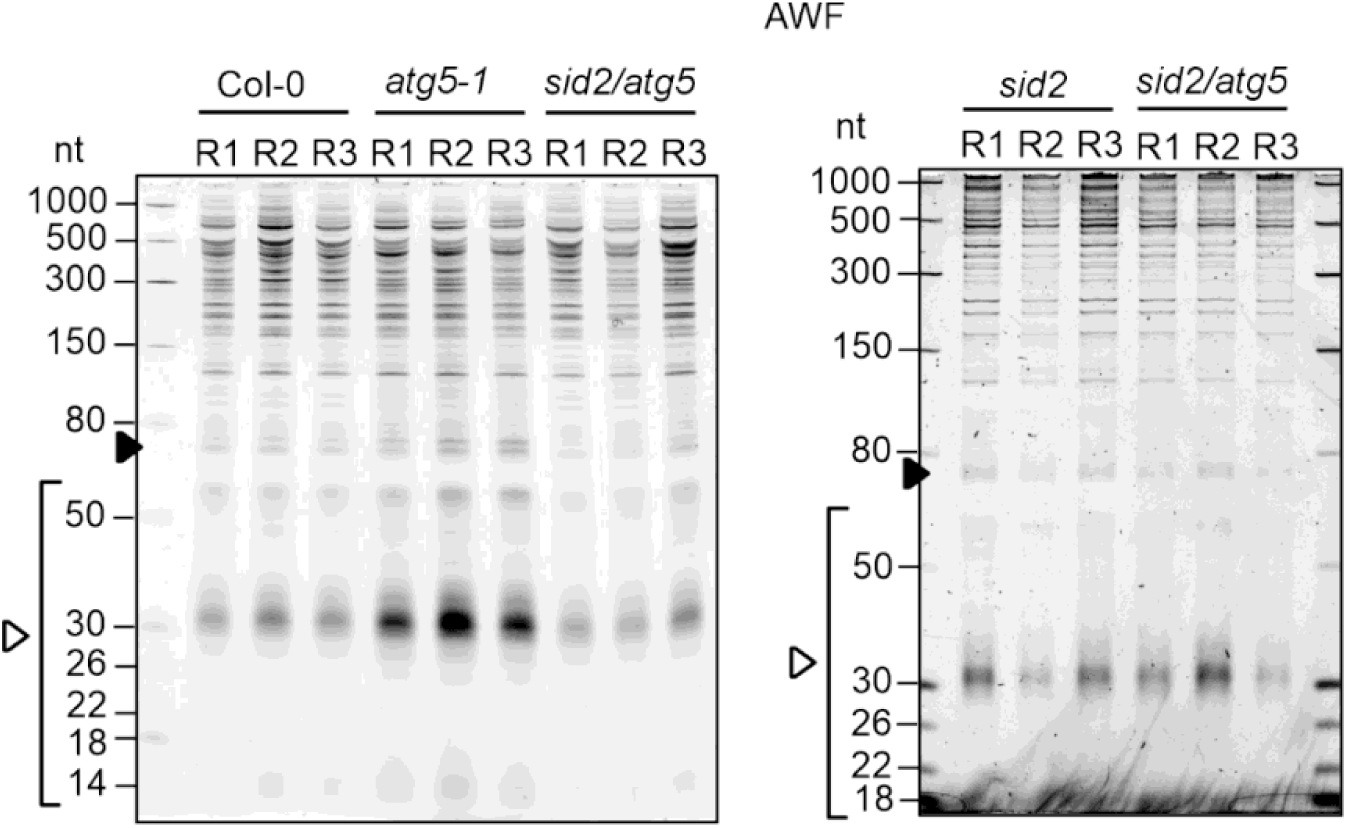
Autophagy is not involved in RNA secretion into the apoplast. Denaturing PAGE of RNA isolated from AWF of Col-0, *atg5*, *sid2*, and *sid2/atg5* plants. Black arrowheads indicate slightly truncated tRNAs, and white arrowheads indicate tRNA fragments (tRFs). RNA was isolated from equal volumes of AWF, corresponding to equivalent plant fresh weights. Three biological replicates (R1–R3) are shown.

**Fig. S6.**
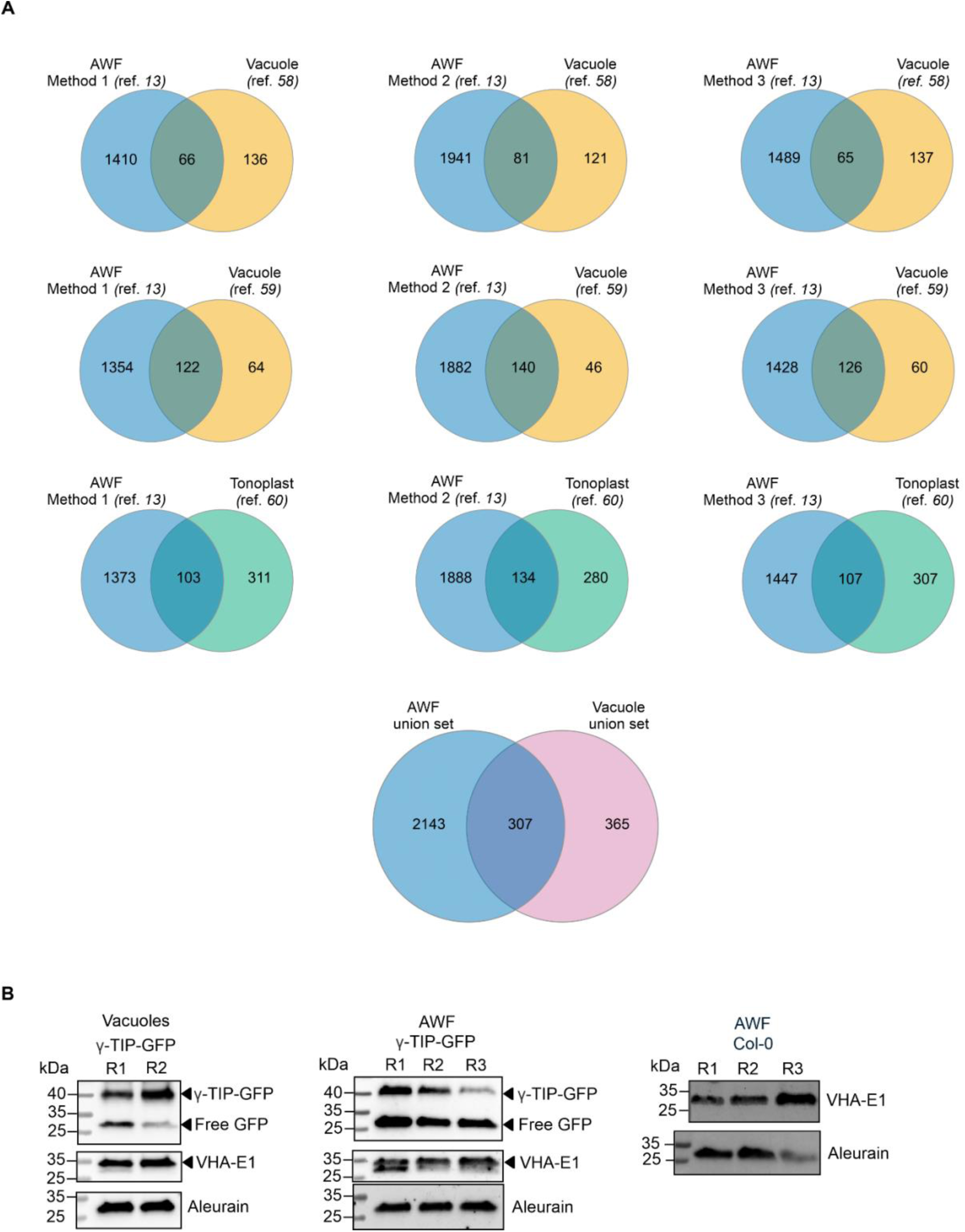
Approximately half of all vacuolar proteins are found in the AWF. **(A)** Venn diagrams showing the overlap between *Arabidopsis* AWF and vacuolar proteomes. Three AWF proteomes (blue) were obtained from (*13*), each generated using a different isolation method as described in the corresponding citation. Vacuolar soluble proteins (orange groups) were derived from vacuoles purified from mature leaves (proteome published in (*58*)) and from intact vacuoles isolated from *Arabidopsis* suspension-cultured cells (proteome published in (*59*)). The tonoplast proteome (green) was obtained from vacuoles isolated from 5 day-old seedlings (proteome published in (*60*). For the union sets, proteins detected in at least one dataset within the same fraction were included. The vacuolar union set (pink) integrates both soluble vacuolar and tonoplast proteins. Numbers indicate protein counts per group. Venn diagrams were created with the online tool Interactivenn https://www.interactivenn.net/. **(B)** Immunoblot detection of γ-TIP-GFP and VHA-E1 (tonoplast markers) and aleurain (a vacuolar lumen marker) in unfiltered AWF. Two (R1 and R2) or three (R1-R3) biological replicates are shown for purified vacuoles and AWF, respectively. Purity of isolated vacuoles is presented in fig. S7.

**Fig. S7.**
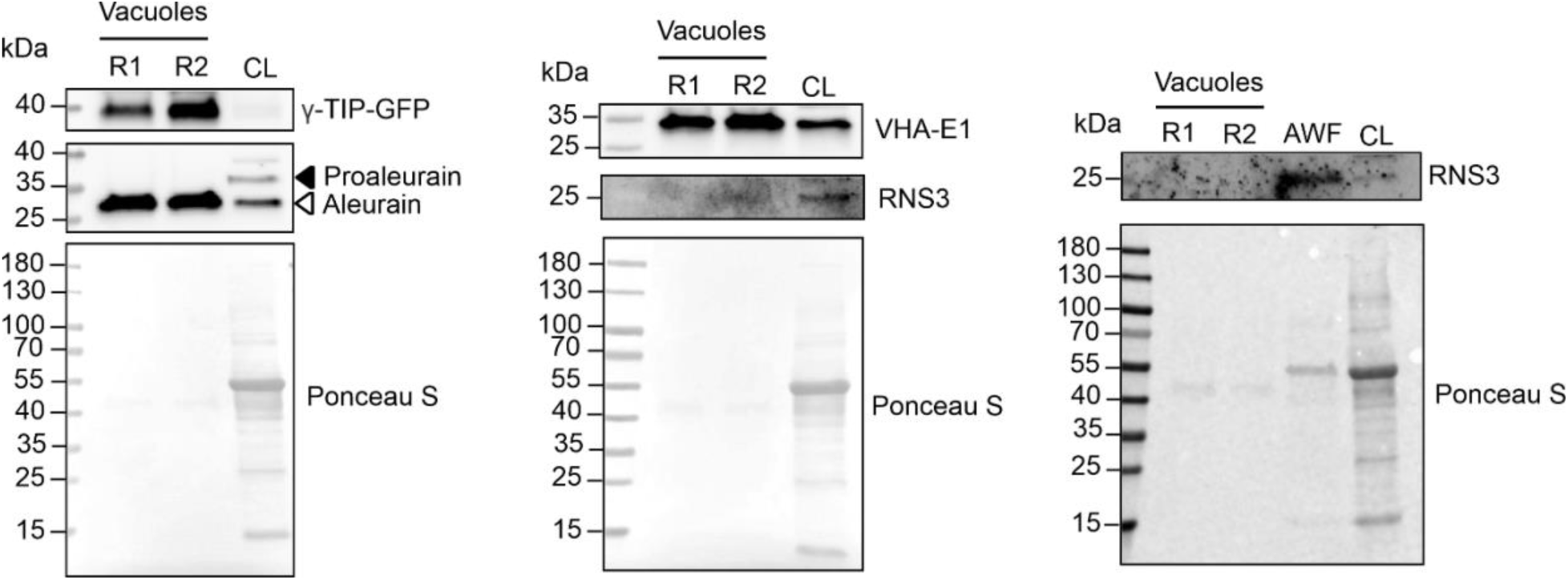
Vacuolar fractions are enriched for vacuolar markers and depleted of apoplastic proteins. Immunoblots detecting the vacuolar proteins aleurain, VHA-E1, and γ-TIP-GFP, and the apoplastic marker RNS3 in cell lysate (CL), AWF, and isolated vacuoles. Protein loading was adjusted for each fraction to enable detection of their respective markers. Similar protein amounts were loaded for AWF and vacuolar fractions, and a higher protein amount was loaded for CL to allow detection of compartment-specific markers that are diluted in the total cellular context.

**Fig. S8.**
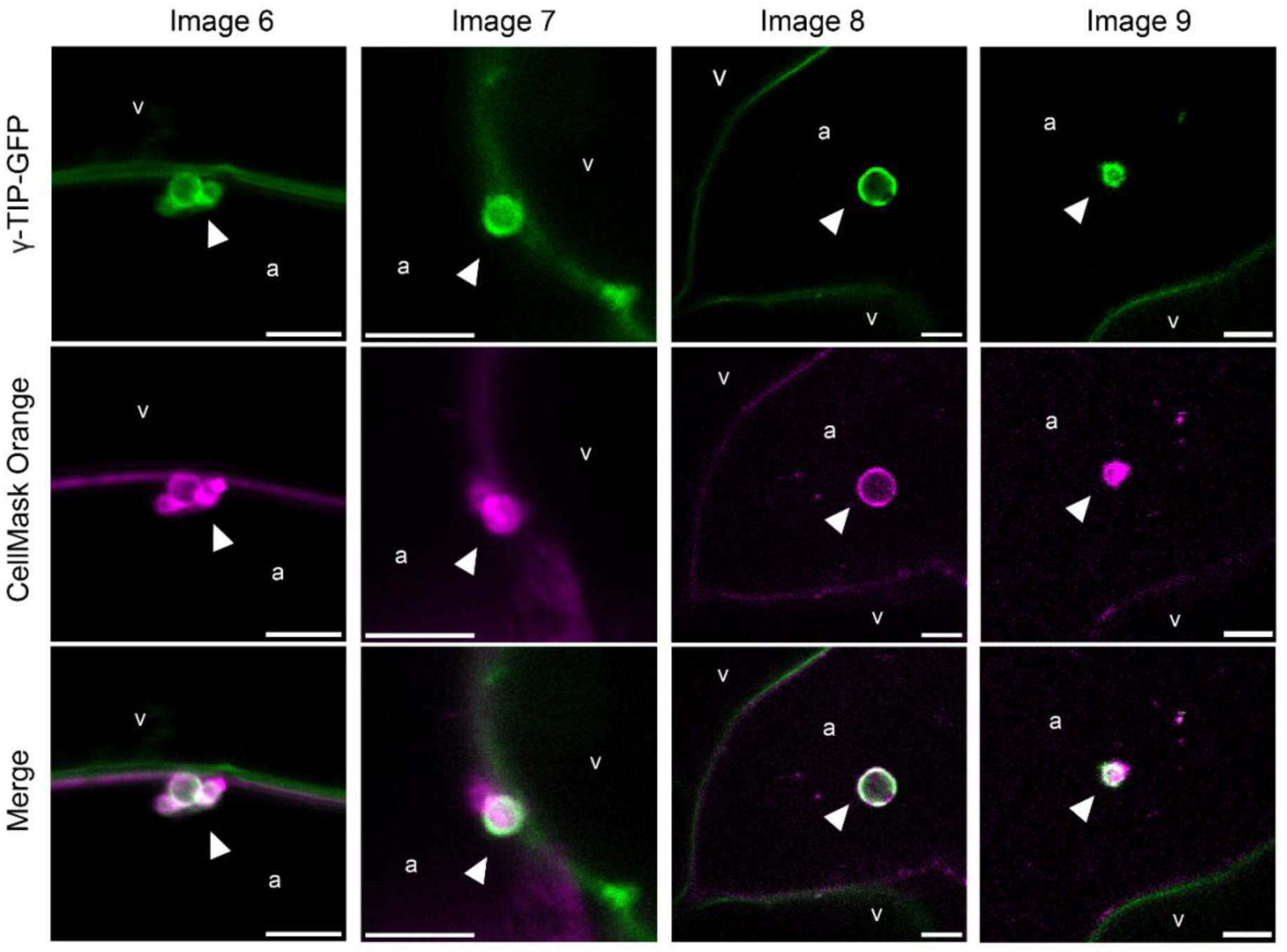
Extracellular Vacuole-derived vesicles (EVacs) containing γ-TIP-GFP are released into the apoplast. In vivo confocal imaging of mesophyll cells from freshly detached leaves of 6-week-old γ-TIP-GFP *Arabidopsis* plants. Plasma membranes were stained with CellMask Orange. To enable imaging of the apoplastic space, leaves were infiltrated with VIB or distilled water and mounted in VIB buffer immediately prior to imaging. Images were acquired on a Leica Stellaris 8 confocal microscope. Vacuoles (v) and apoplast (a) are indicated. Arrowheads mark EVacs. Scale bars, 5 µm. See Movies S1-S6 for additional EVac examples.

**Fig. S9.**
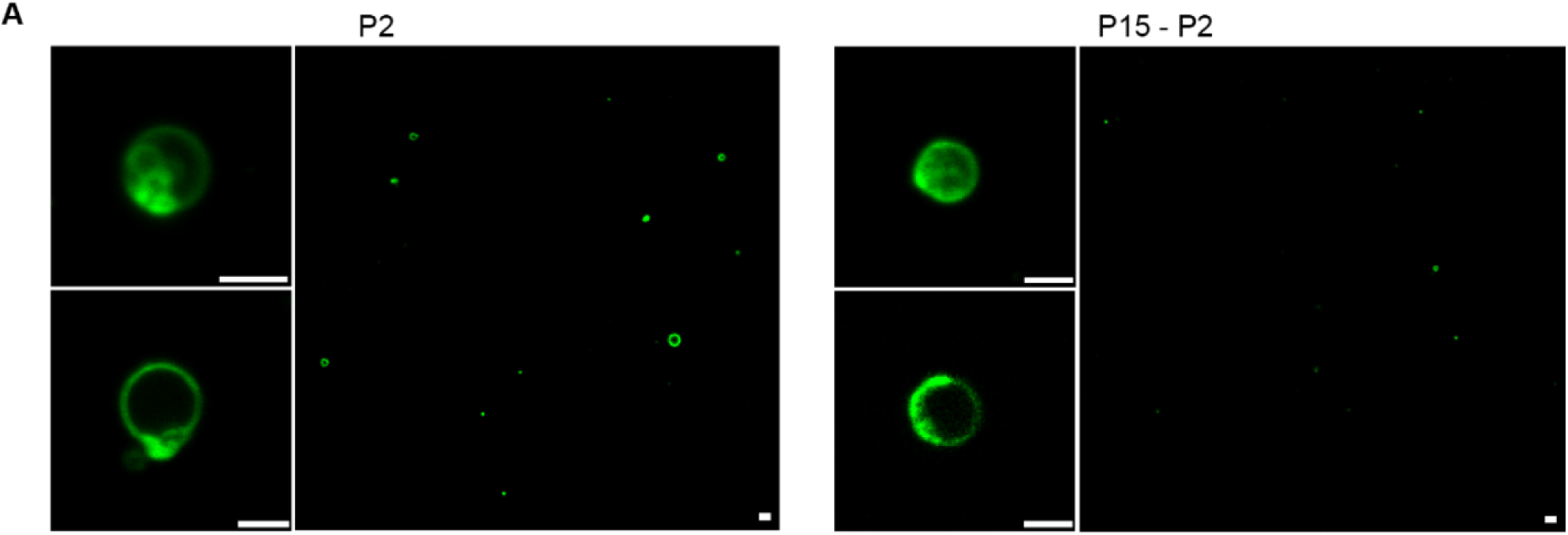
P2 and P15-P2 fractions contain EVacs. Confocal images of P2 and P15-P2 fractions showing enrichment of EVacs. For each fraction, an average intensity projection (top left panel) and single optical sections (bottom left panel and right panel) are shown. Scale bar, 2 µm.

**Fig. S10.**
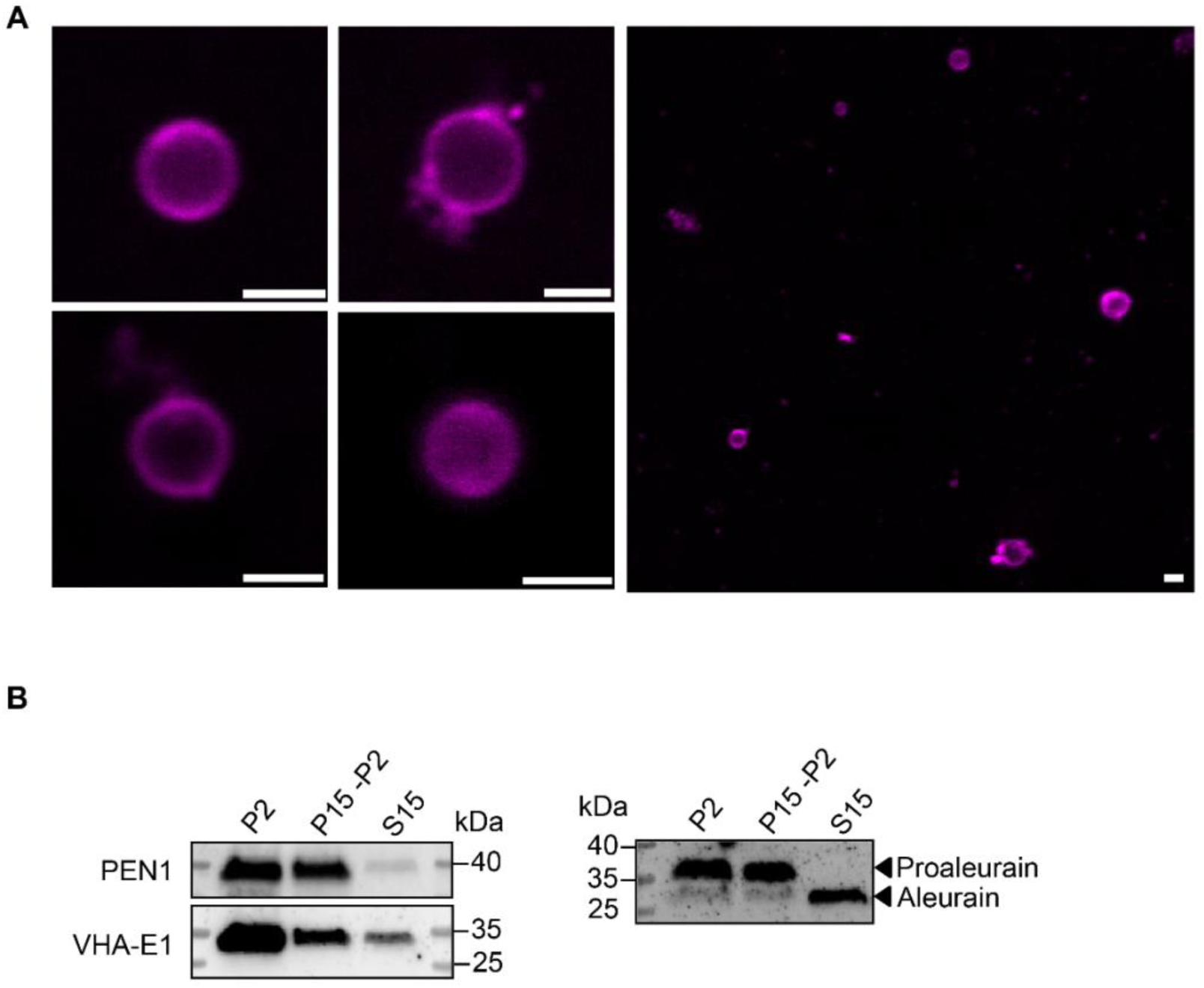
EVacs can also be isolated from non-transgenic plants. **(A)** Confocal images of EVacs (P10) isolated from wild-type Col-0 plants. EVac membranes were labeled with CellMask Orange. Scale bar, 2 µm. **(B)** Immunoblots showing enrichment of vacuolar markers in EVac-containing fractions isolated from wild-type Col-0 plants. Loading is described in Materials and Methods.

**Fig. S11.**
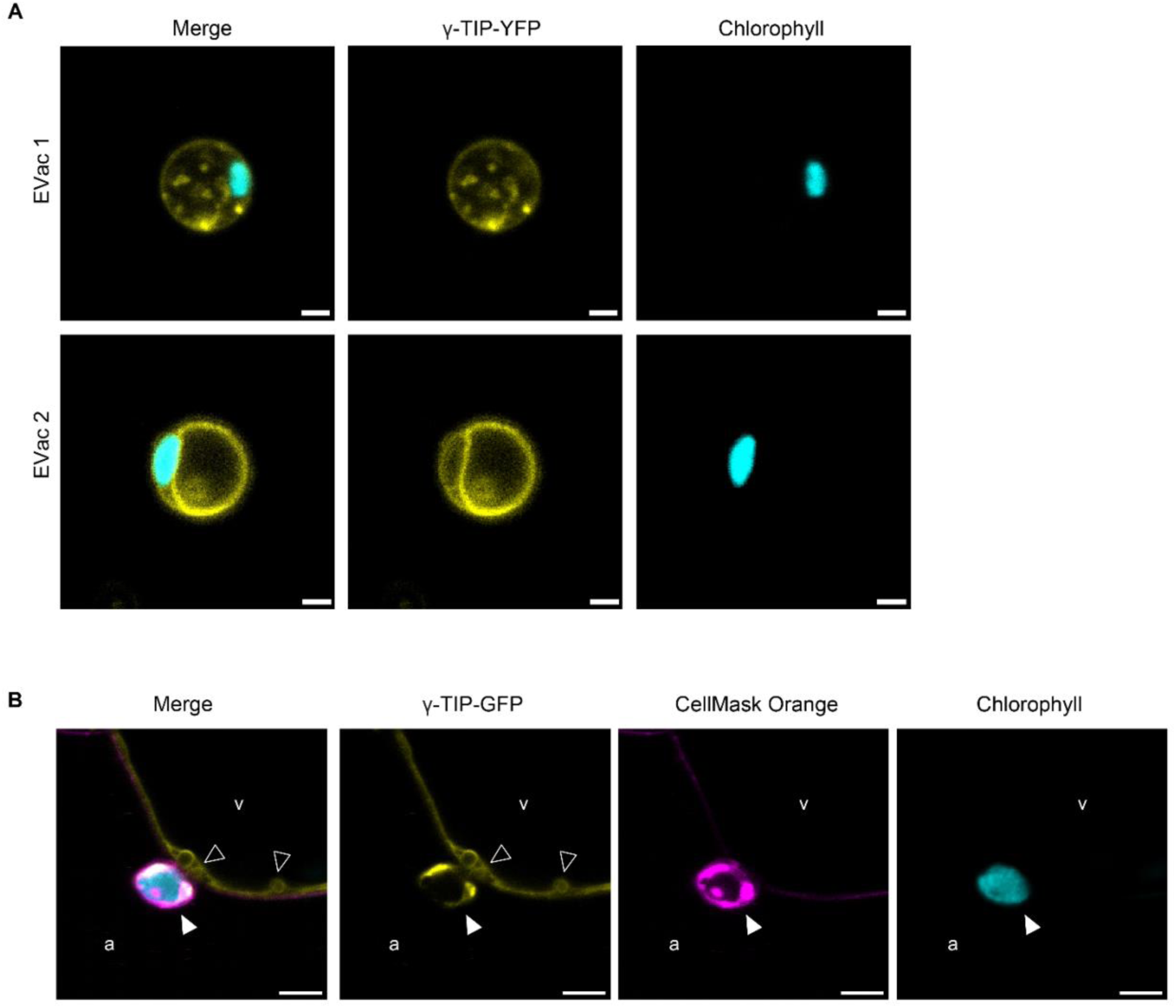
Chloroplasts can be secreted inside EVacs. **(A)** Confocal images of EVacs (P10) isolated from the γ-TIP-YFP line, showing chloroplasts inside the EVacs. Scale bar, 2 µm. **(B)** In vivo confocal imaging of mesophyll cells from 6-week-old γ-TIP-GFP *Arabidopsis* leaves. Plasma membranes were stained with CellMask Orange. Leaves were infiltrated with VIB or distilled water prior to imaging. Images were acquired on a Leica Stellaris 8 confocal microscope. Vacuoles (v) and apoplast (a) are indicated. Filled arrowhead marks an EVac encapsulating a chloroplast and open arrowheads mark vacuole-derived bodies in the cytoplasm. Scale bar, 5 µm.

**Fig. S12.**
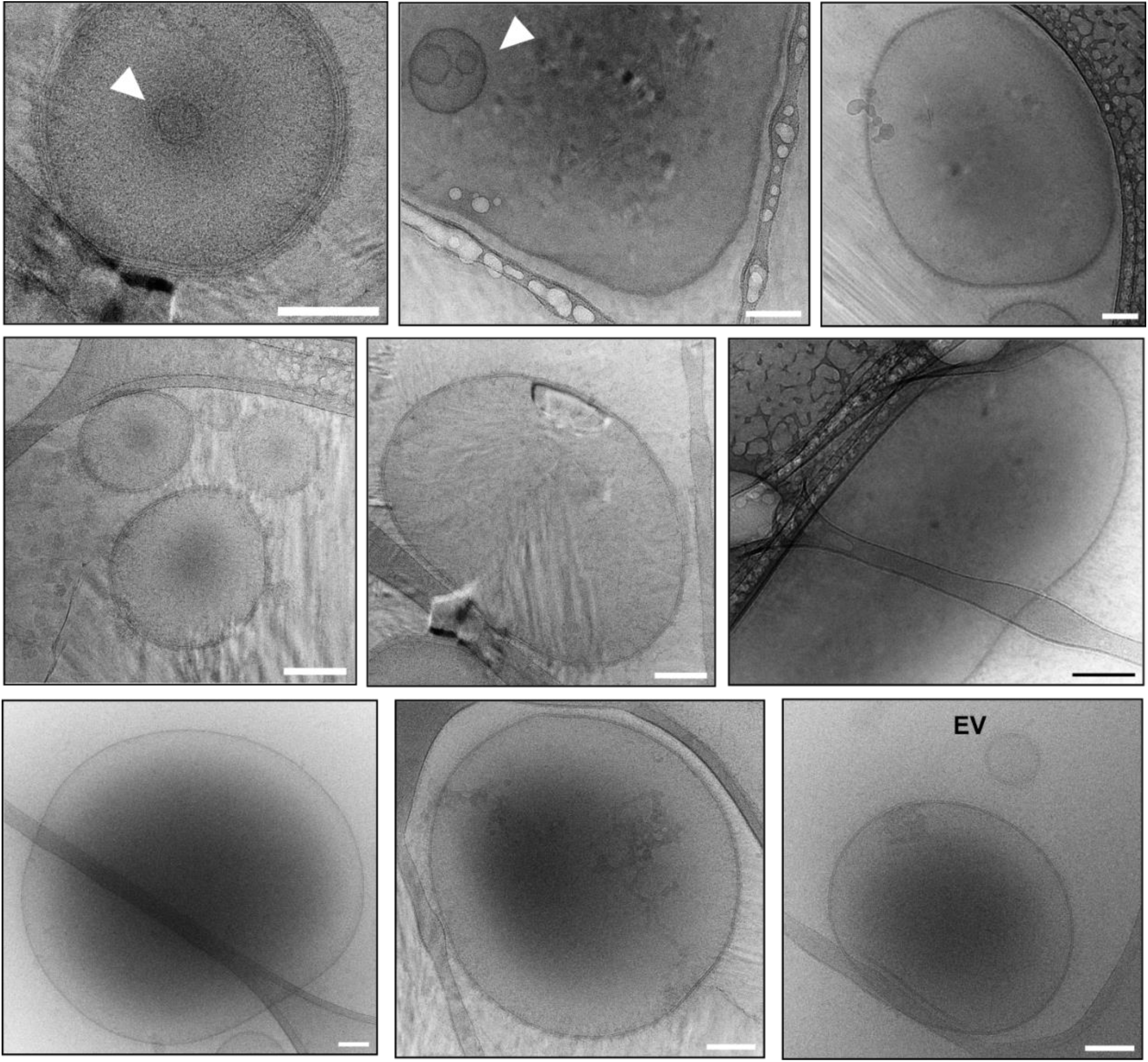
Morphological diversity of EVacs. Representative cryo-EM images of EVacs in a P10 fraction, showing the range of sizes and structural features observed, including single and double membrane delimitation and the presence of intraluminal vesicles (arrowheads). The bottom right image shows a conventional extracellular vesicle (EV) alongside an EVac for size comparison. Scale bars, 100 nm.

**Fig. S13.**
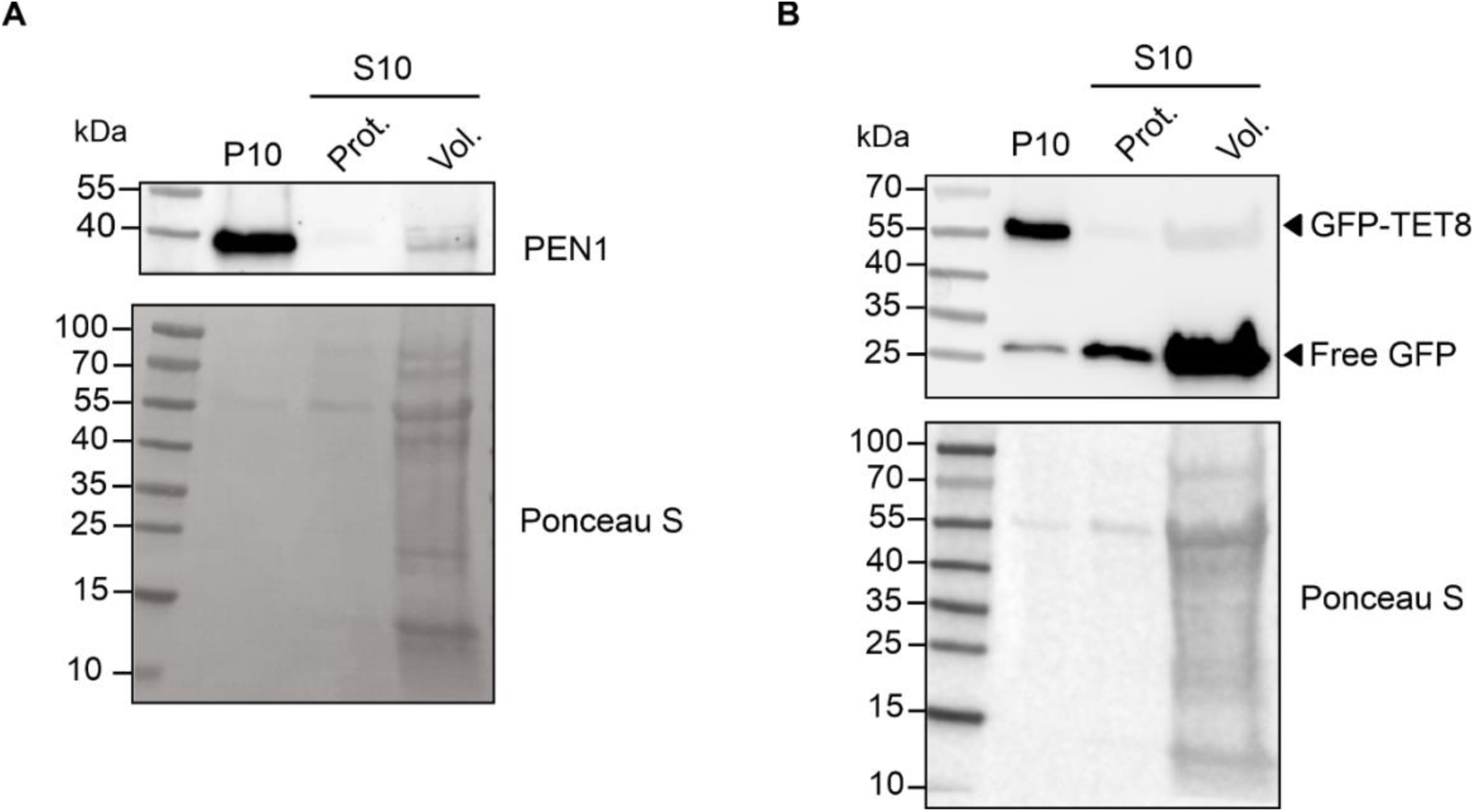
The EV markers PEN1 and TET8 are enriched in P10 fractions. **(A)** Immunoblot detecting the EV marker PEN1 in P10 and S10 fractions. Blot was probed with anti-PEN1 antibody. Lanes represent the P10 fraction, the S10 fraction loaded at an equal protein amount (Prot.) relative to P10, and the S10 fraction corresponding to the same original volume of AWF (Vol.) used for the P10 preparation. The S10 normalized per volume was prepared by acetone precipitation of the total supernatant remaining after P10 collection. **(B)** Immunoblot detecting the EV marker GFP-TET8 in P10 and S10 fractions. Blot was probed with anti-GFP antibody. For this blot, a GFP-TET8 line was used for P10 and S10 collection. Loading was done as in panel A.

**Fig. S14.**
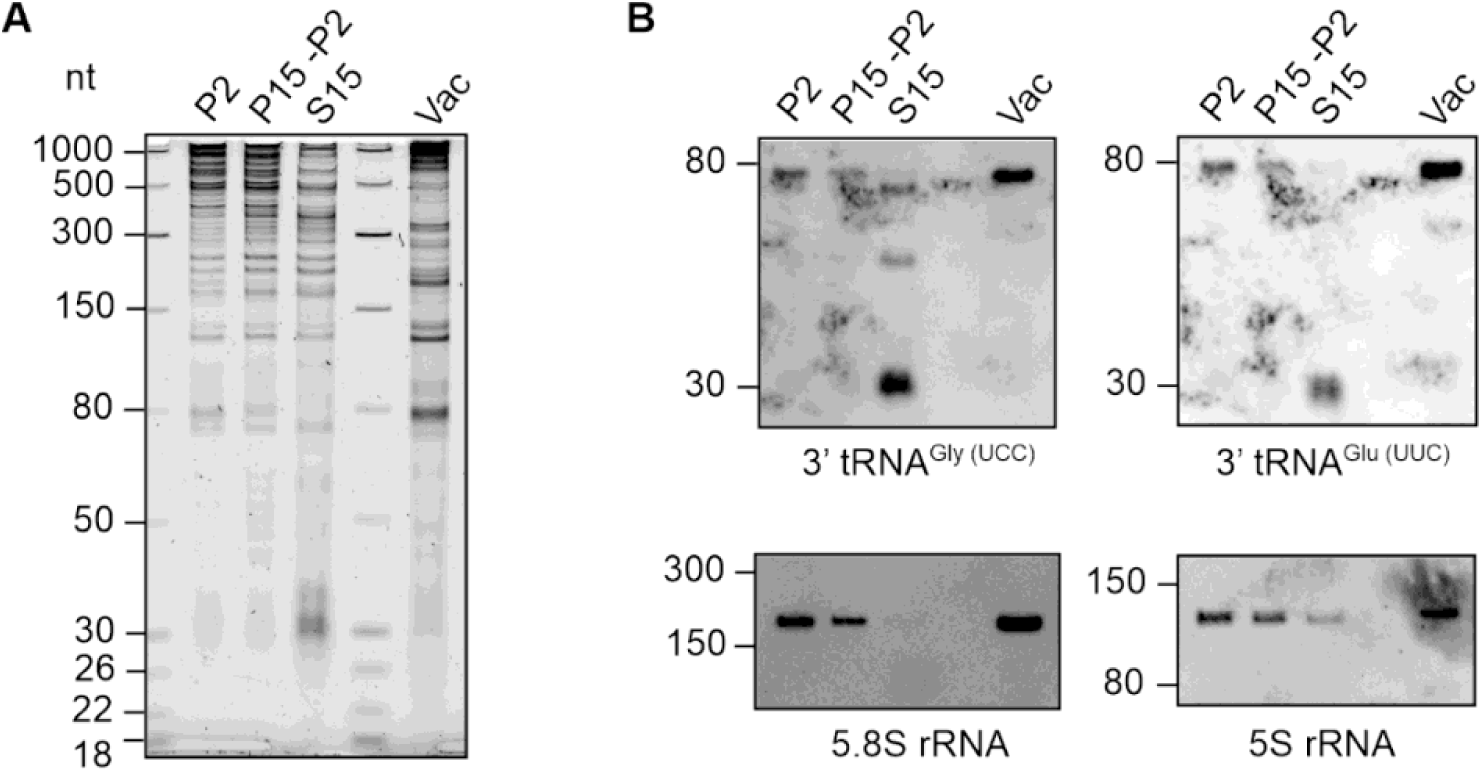
EVacs isolated from wild-type Col-0 plants show an RNA pattern similar to EVacs from γ-TIP-GFP transgenic plants. **(A)** Denaturing PAGE showing the RNA pattern of P2, P15-P2 and S15 as well as vacuoles isolated from wild-type Col-0 plants. **(B)** Northern blot detection of the indicated RNAs from the gel shown in (A).

**Fig. S15.**
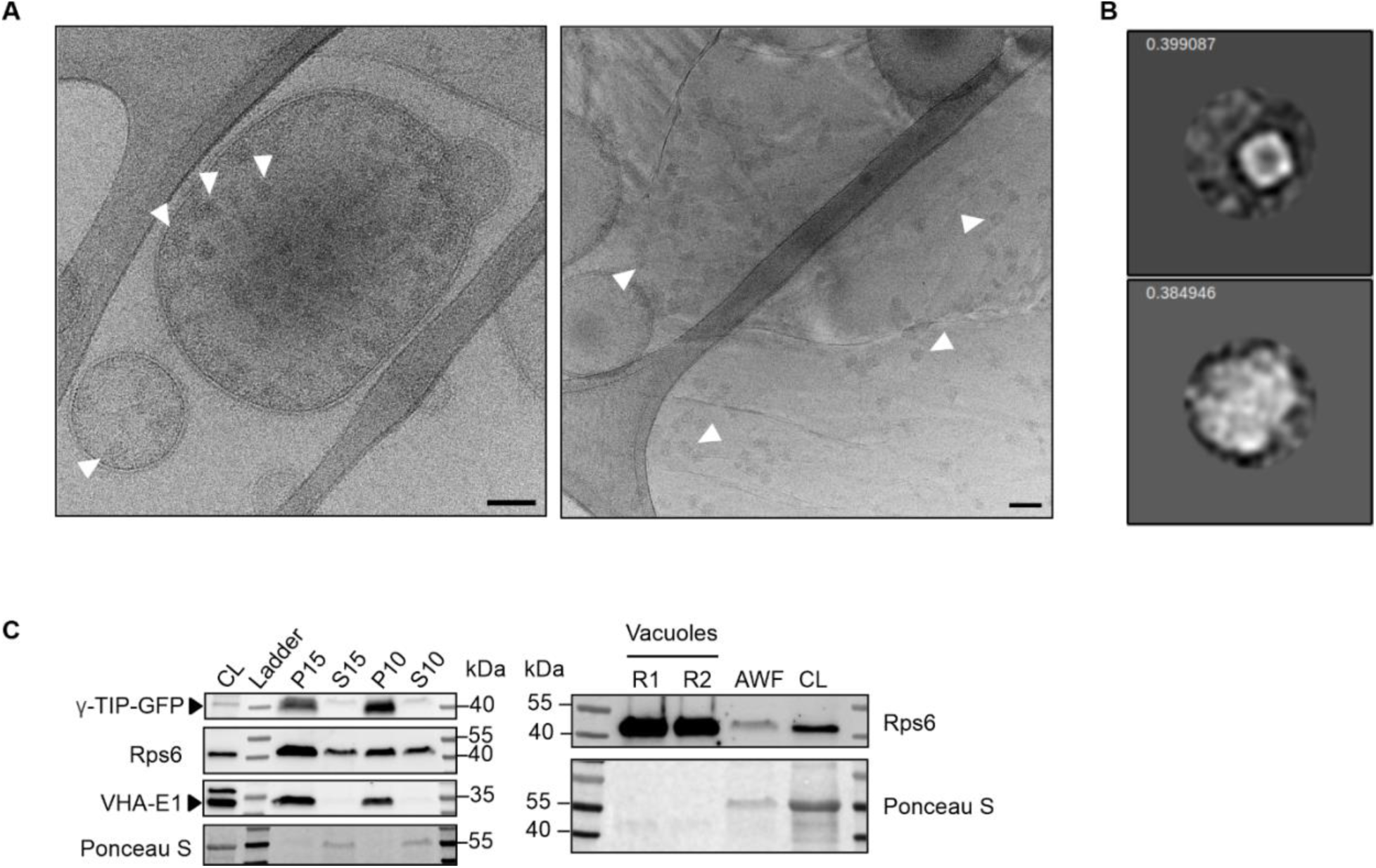
EVac fractions contain ribosomes. **(A)** Cryo-EM images of purified EVac fractions showing ribosome-like particles inside EVacs (arrowheads). Scale bars, 50 nm. **(B)** 2D class average of two size populations for particles in the EVac fraction. The small and large classes contain 85 and 82 particles, respectively. The top left values show class distribution in the final classification, which used 213 particles cleaned up from 573 initial picks. The spherical mask diameter and the box size are 300 Å and 576.24 Å, respectively**. (C)** Left, immunoblots showing enrichment of the small ribosomal subunit protein Rps6 and the tonoplast marker VHA-E1 in EVac-containing pellets (P15 and P10) relative to their corresponding supernatants (S15 and S10). Arrowheads indicate the expected band for each protein. Loading is described in Materials and Methods. Right, enrichment of Rps6 in isolated vacuoles relative to AWF and total cell lysate (CL). Loading is described in Materials and Methods.

**Fig. S16.**
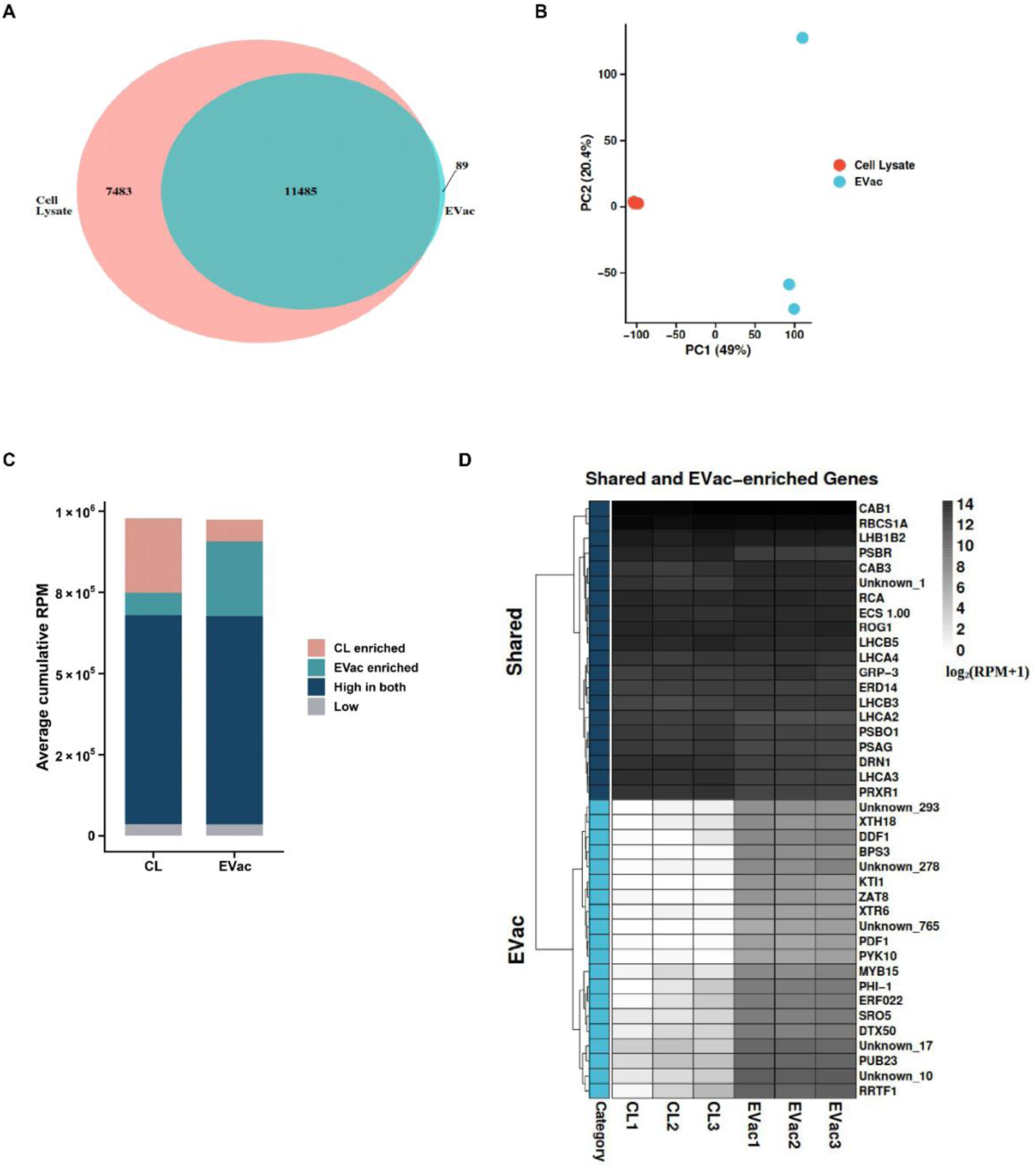
EVacs contain mRNAs representative of total cellular RNA. **(A)** Venn diagram showing overlap between mRNAs identified in EVac RNA compared to total cell lysate RNA. Genes were considered expressed if they had ≥10 raw read counts in at least two of the three biological replicates within each group. The diagram illustrates the numbers of genes uniquely detected in each group and those shared between both groups. **(B)** Principal component analysis (PCA) of 3′ RNA-seq data was performed using RPM (reads per million mapped reads) values. Each point represents an individual biological replicate, colored by experimental group (Orange: Cell Lysate; Blue: EVac). The percentages on the axes indicate the variance explained by each principal component. **(C)** Distribution of cumulative transcript abundance in total cell lysate and EVac samples. Genes were classified based on mean expression - into cell lysate-enriched, EVac-enriched, high in both (shared), or low-expression categories (RPM ≤10). Stacked bars represent the average cumulative RPM contributed by each category. **(D)** Heatmap of shared and EVac-enriched transcripts. The top 20 shared and top 20 EVac-enriched mRNAs are shown based on log₂ (RPM + 1) across biological replicates. Darker shading indicates higher RPM.

**Fig. S17.**
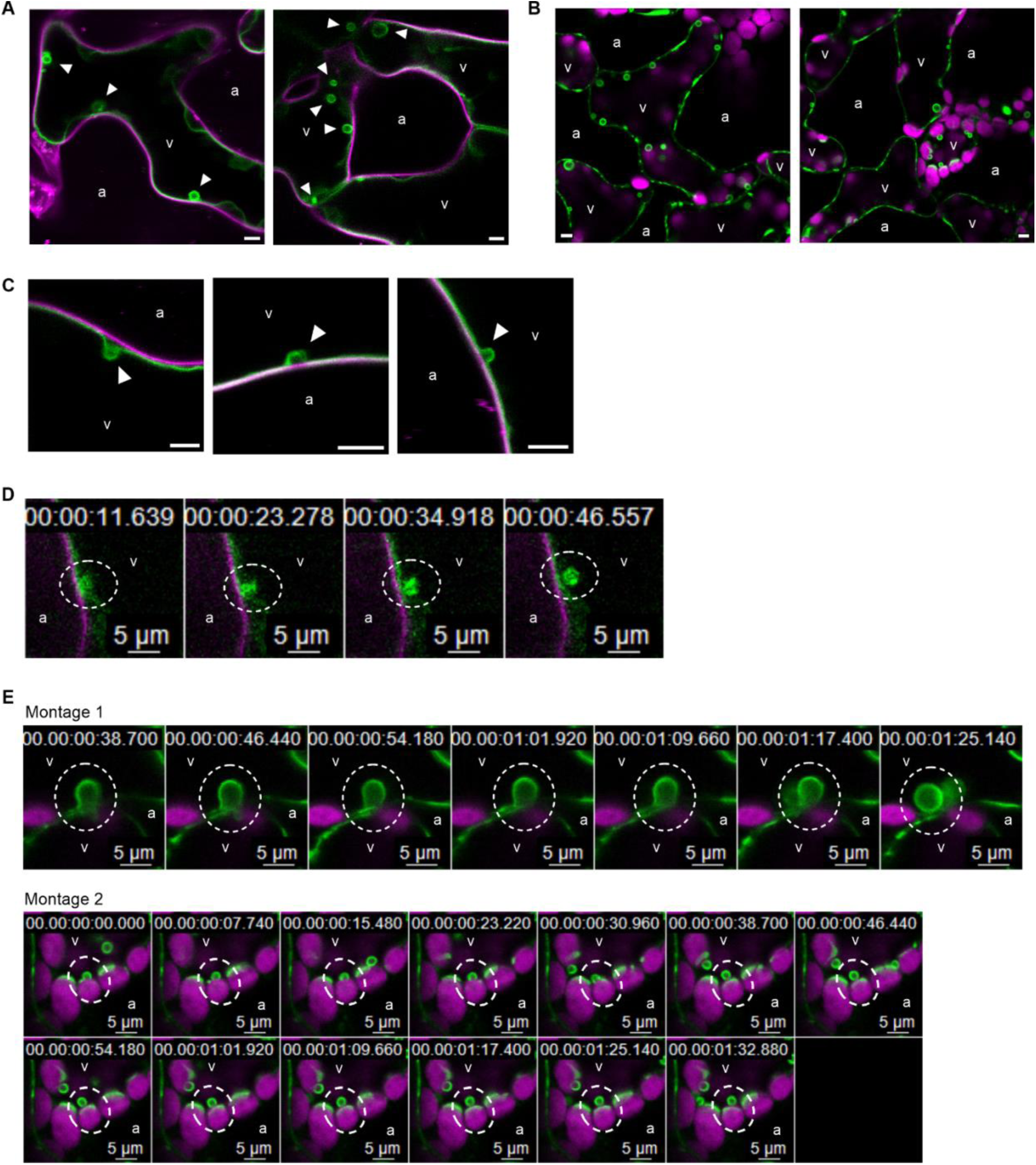
Intravacuolar body biogenesis via tonoplast inward folding in mesophyll cells. In vivo confocal imaging of mesophyll cells from freshly detached leaves. γ-TIP-GFP/YFP signal is shown in green throughout. Vacuoles (v) and apoplast (a) are indicated. All scale bars, 5 µm. **(A, B)** Single optical sections showing intravacuolar bodies inside vacuoles**. (A)** Mesophyll cells from 6-week-old γ-TIP-GFP plants. Plasma membrane labeled with CellMask Orange is shown in magenta. Arrowheads indicate intravacuolar bodies. **(B)** Mesophyll cells from 2-week-old γ-TIP-YFP seedlings. Chloroplast autofluorescence is shown in magenta. **(C)** Single optical sections from γ-TIP-GFP leaves illustrating tonoplast invagination during intravacuolar body formation. Arrowheads indicate the site of invagination. Plasma membrane stained with CellMask Orange is shown in magenta. **(D–E)** Time-lapse montages showing intravacuolar body formation via tonoplast invagination. Dashed circles indicate the forming intravacuolar body. **(D)** Mesophyll cells from 6-week-old γ-TIP-GFP plants. Plasma membrane stained with CellMask Orange is shown in magenta. **(E)** Mesophyll cells from 2-week-old γ-TIP-YFP seedlings. Chloroplast autofluorescence is shown in magenta. Two independent time-lapse series (Montage 1 and Montage 2) are shown.

**Fig. S18.**
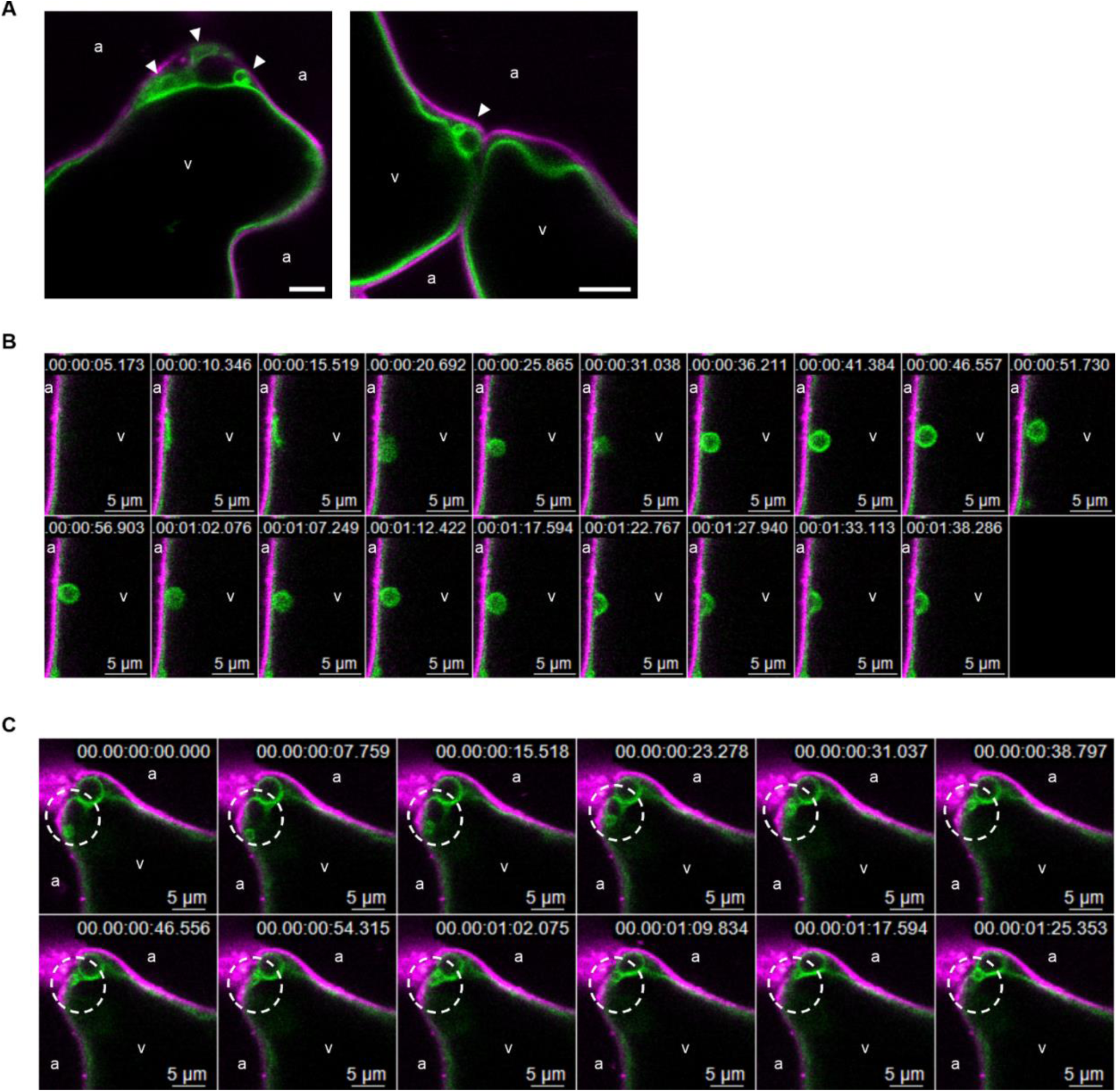
Vacuole-derived bodies are released from the vacuole into the cytosol. In vivo confocal imaging of mesophyll cells from freshly detached leaves. γ-TIP-GFP is shown in green; plasma membrane, stained with CellMask Orange, is shown in magenta. Vacuoles (v) and apoplast (a) are indicated. All scale bars, 5 µm. **(A)** Single optical sections showing vacuole-derived bodies (arrowheads) located in the cytosol of mesophyll cells from two independent γ-TIP-GFP plants. **(B)** Time-lapse montage showing an intravacuolar body forming via tonoplast invagination and subsequently being released into the cytosol. **(C)** Time-lapse montage showing a vacuole-derived body (dashed circles) forming directly in the cytosol via tonoplast folding.

**Fig. S19.**
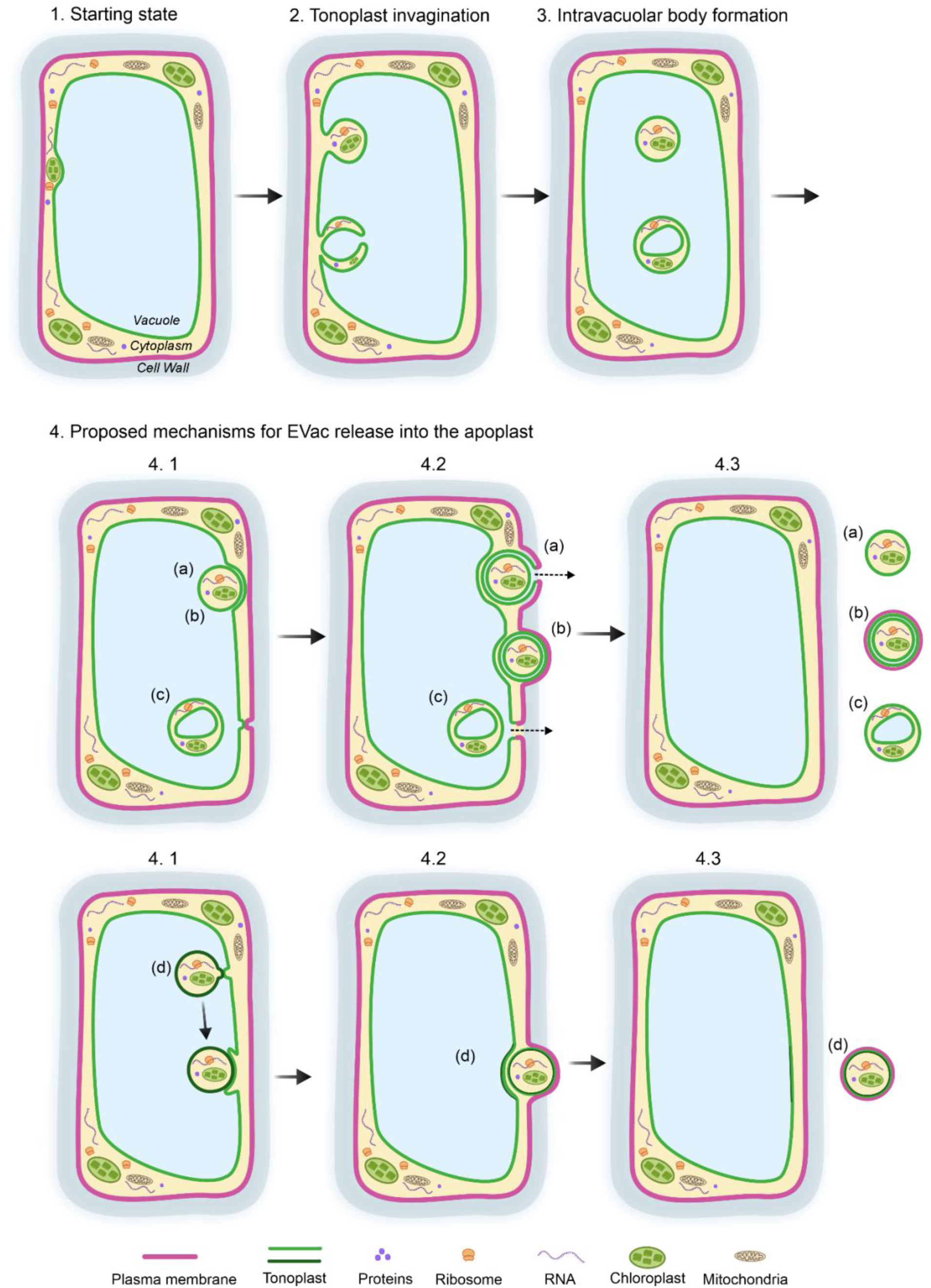
Proposed model for EVac secretion into the extracellular space. Several biogenesis pathways for intravacuolar bodies have been previously proposed (*21, 29*). Our confocal time-lapse imaging suggests that intravacuolar bodies are formed via tonoplast invagination and folding, which engulfs cytoplasmic content (2). This process can generate either a single-membrane intravacuolar body containing only cytosolic cargo (3, top) or a double-membrane body enclosing vacuolar cargo as well cytosol content between the two membrane layers (3, bottom). These intravacuolar bodies can subsequently escape the vacuole and be released to the apoplast (4). Our data suggests that multiple mechanisms might be involved. We propose four possibilities. In mechanisms (a) and (b), the EVac buds off from the vacuole (4.1) and transits the cytoplasm as a double-membrane body (4.2). Subsequently, the outer membrane of the double-membrane body can either fuse with the plasma membrane, releasing the internal EVac as a single-lipid bilayer vesicle (a, 4.3), or the double-membrane cytosolic body can be released into the apoplast by a subsequent budding mechanism that yields a multilayer EVac (b, 4.3). In mechanism (c), an intravacuolar body can directly translocate from the vacuole to the apoplast through a localized direct fusion event between the tonoplast and the plasma membrane. In mechanism (d), intravacuolar bodies escape the vacuole via a hemifusion-mediated budding mechanism (*36*), releasing a single bilayer cytosolic body, that subsequently can reach the apoplast as a double-membrane EVac via a budding mechanism.

**Table S1.**
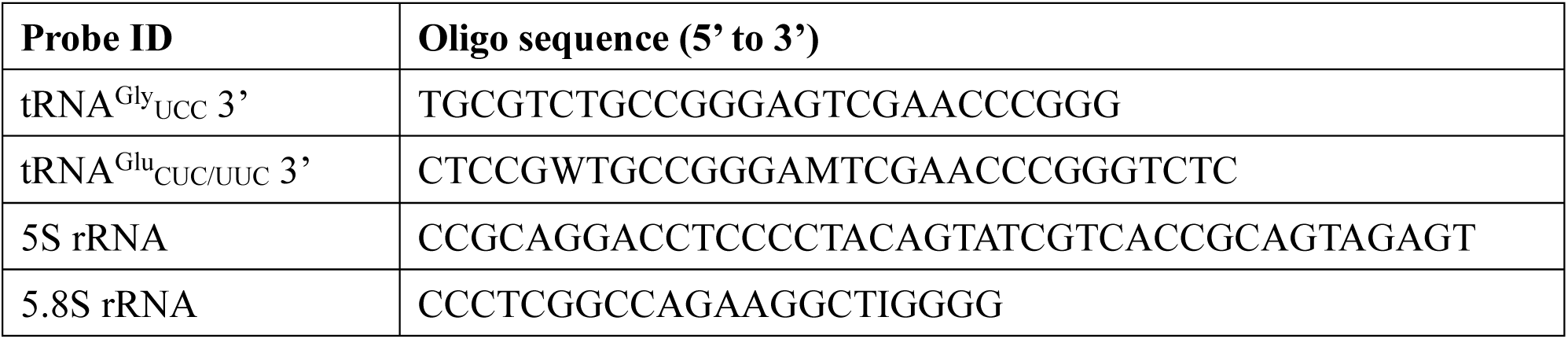
Oligonucleotide sequences of hybridization probes.

**Movie S1.** Budding and release of an EVac into the apoplastic space In vivo confocal time-lapse sequence of mesophyll cells from freshly detached leaves of a 6-week-old *Arabidopsis* plant expressing γ-TIP-GFP. The apoplastic space was infiltrated with VIB buffer or water prior to imaging. Vacuolar membranes (tonoplast) are labeled with γ-TIP-GFP (green) and the plasma membrane is stained with CellMask Orange (magenta). The sequence captures an EVac actively budding off from the plasma membrane and another EVac floating in the apoplast. Scale bar, 5 µm.

**Movie S2.** EVac floating dynamically within the apoplastic space in young plants In vivo confocal time-lapse sequence of mesophyll cells from freshly detached leaves of a 2-week-old *Arabidopsis* seedling expressing γ-TIP-YFP. The apoplastic space was infiltrated with VIB buffer or water prior to imaging. Vacuolar membranes (tonoplast) are labeled with γ-TIP-YFP (green) and chloroplasts are shown in magenta. Scale bar, 5 µm.

**Movie S3.** EVacs floating dynamically within the apoplastic space in young plants In vivo confocal time-lapse sequence of mesophyll cells from freshly detached leaves of a 2-week-old *Arabidopsis* seedling. The plant is dual-labeled, expressing γ-TIP-GFP (green) to mark vacuolar membranes (tonoplast) and mCherry-SYP61 (magenta) as a trans-Golgi network (TGN) marker. The apoplastic space was infiltrated with VIB buffer or water prior to imaging. Scale bar, 5 µm.

**Movies S4-S6.** EVacs floating dynamically within the apoplastic space in mature plants In vivo confocal time-lapse sequences of mesophyll cells from freshly detached leaves of a 6-week-old *Arabidopsis* plant expressing γ-TIP-GFP. The apoplastic space was infiltrated with VIB buffer or water prior to imaging. Vacuolar membranes (tonoplast) are labeled with γ-TIP-GFP (green) and the plasma membrane is stained with CellMask Orange (magenta). The movies show different examples of EVacs floating in the apoplast. Scale bar, 5 µm.

**Movie S7.** Co-localization of γ-TIP-GFP and RFP-PEN1 in an Evac In vivo confocal time-lapse sequence of mesophyll cells from freshly detached leaves of a dual-labeled 2-week-old *Arabidopsis* seedling expressing γ-TIP-GFP (in green) and RFP-PEN1 (in magenta). The apoplastic space was infiltrated with VIB buffer or water prior to imaging. The video shows a dual-labeled EVac floating in the apoplast. In this Evac, both γ-TIP-GFP and RFP-PEN1 co-localize at the EVac membrane. Scale bar, 5 µm.

**Movie S8.** Vacuole-derived body moving in the cytosol In vivo confocal time-lapse sequences of mesophyll cells from freshly detached leaves of a 6-week-old *Arabidopsis* plant expressing γ-TIP-GFP. The apoplastic space was infiltrated with VIB buffer or water prior to imaging. Vacuolar membranes (tonoplast) are labeled with γ-TIP-GFP (green) and the plasma membrane is stained with CellMask Orange(magenta). The video captures a vacuole-derived body that has escaped the vacuole and moves in the cytosol. Scale bar, 5 µm.

**Data S1.** (separate file) High-confidence proteins identified by LC–MS/MS following protein-level false discovery rate (FDR) filtering.

**Data S2.** (separate file) 3’ polyA RNA-seq analysis summary. Separate worksheets contain sequencing quality control, read mapping statistics, normalized read counts (RPM values), and the top 20 EVac enriched and shared candidate genes for three biological replicates.

